# Probing the structural heterogeneity of Pup ligase PafA using H/D exchange mass spectrometry

**DOI:** 10.1101/2024.12.16.628271

**Authors:** Alicia Plourde, Jacquelyn C. Ogata-Bean, Siavash Vahidi

## Abstract

The Pup-proteasome system (PPS) is a unique bacterial proteolytic pathway found in some bacterial species, including in *Mycobacterium tuberculosis*, that plays a vital role in maintaining proteome integrity and survival during infection. Pupylation is the process of tagging substrates with Pup for degradation and is catalyzed by PafA, the sole Pup ligase in bacteria. However, how PafA interacts with diverse targets and its oligomeric state remain poorly understood. Although X-ray crystal structures have characterized PafA as a domain-swapped dimer, it is widely regarded as functionally active in its monomeric form. It remains to be established whether PafA dimerizes in solution, and how dimerization influences its function. In this study, we employed hydrogen-deuterium exchange mass spectrometry (HDX-MS) alongside complementary biophysical techniques to explore the oligomeric states and conformational dynamics of PafA. We show that recombinantly-produced PafA exists in a monomeric and a domain-swapped dimeric state in solution. While PafA_dimer_ is stabilized by nucleotide, it predominately resides in a catalytically inactive conformation. Our HDX-MS highlighted regions throughout the N- and C-terminal domains that facilitate the PafA dimerization process. HDX-MS also revealed nucleotide binding induces global conformational changes on the PafA_monomer_, underscoring the structural plasticity of this promiscuous enzyme. Our findings provide insight into the structure-function-dynamics relationship of PafA, shedding light on how its oligomeric states and conformational flexibility might influence its role in the PPS.

## INTRODUCTION

Members of the Actinobacteria and Nitrospirota phyla, including pathogenic mycobacteria, encode a rare Pup-proteasome system (PPS) (1). This system uses the intrinsically disordered prokaryotic ubiquitin-like protein (Pup) to tag proteins for proteasomal degradation via a process called pupylation (1, 2). The PPS contributes to proteome integrity and is vital for *Mycobacterium tuberculosis* survival during infection (1, 3–5). Pupylation is performed by a pair of structurally homologous enzymes, Dop (deaminase of Pup) and PafA (proteasome accessory factor A). Dop deaminates Pup’s C-terminal glutamine into glutamate, thereby making it ligation competent (**Fig. S1A**) (6). Subsequently, PafA catalyzes an ATP-dependent two-step reaction to ligate Pup to target proteins by forming a phosphorylated Pup intermediate and subsequent ligation of Pup to a lysine residue of the target protein (**Fig S1B**). Pupylated proteins are either recognized and degraded by the proteasome system or spared through Dop’s depupylase activity (**Fig S1C**) (5, 7–9). The interplay between PafA-mediated pupylation and Dop- mediated depupylation is thought to determine the cellular pupylome.

Despite the functional similarities between the substrate tagging mechanism of the PPS and ubiquitin-proteasome system (UPS), the enzymes involved in these pathways are distinct and lack structural homology (6, 10–13). PafA and Dop belong to the carboxylate amine ligase superfamily (14). X-ray crystallographic studies reveal that PafA and Dop adopt similar globular folds with a large N-terminal domain (NTD) that interacts closely with a small C-terminal domain (CTD) (6–8, 15). The NTD shares homology with other carboxylate amine ligases and features a central twisted β-sheet cradle (**Fig. S1D- F**) (6, 14, 16). The concave face of this cradle includes the active site (β-strands 1-4, 6, and 7) and the convex side is protected by bundles of α-helices (6, 15). Interestingly, the CTD is a unique feature of the PafA/Dop family and is absent in other members of the carboxylate amine ligase superfamily (6). Loops in the CTD contribute to the nucleotide-binding pocket, with direct involvement of R418 and W440 that interact with the adenine ring and sugar moiety of the nucleotide, respectively (6).

A notable feature exclusive to Dop is the Dop loop, a conserved loop approximately 40 amino acids in length that precedes β2 (**Fig. S1E** and **Fig. S2**) (6). The Dop loop is located close to the active site and regulates depupylase activity (6, 17–19). Deletion of the Dop loop increases the rate of depupylation *in vitro* and *in vivo* in mycobacteria (19, 20). In all available X-ray crystallographic structures, Dop consistently crystallizes as a monomer (6–8) thus it likely functions in a monomeric state. A recent electron cryomicroscopy (cryo-EM) structure shows monomeric Dop bound to Pup and a hexameric depupylation regulator, with insufficient density to model the Dop loop and Dop’s CTD, suggesting that these regions are inherently dynamic (21). PafA has two reported crystal structures. The first crystal structure [PDB 4B0T (6)] reveals a domain-swapped dimer where N-terminal strand-helix motifs (β1/α1) trade places (coloured blue in **Fig. S1D**), while the active sites face towards one another and the CTDs make numerous secondary contacts with the active site of the other subunit. Intriguingly, the short hinge region, a short segment that connects two swapping domains and undergoes a conformational change during domain swapping, also precedes β2 but is considerably shorter than the Dop loop (**Fig. S2**) (6). In the second PafA crystal structure [PDB 4BJR (15)], a single polypeptide chain containing PafA and a truncated Pup was used (15). Here, the hinge region folds back on itself, forming PafA_monomer_, with Pup from the other chain inserted into the Pup-binding site (15). Despite crystalizing as a domain-swapped dimer, PafA is discussed in the context of a functionally active PafA_monomer_ throughout the literature. The notable exception is the work by Gur and colleagues, who demonstrated that *Mycobacterium smegmatis* PafA forms active dimers in solution and binds target proteins, but not Pup, cooperatively (22). Interestingly, most members of the carboxylate amine ligase superfamily, including glutamine synthetase, adopt higher-order oligomeric structures (6, 16, 23, 24). While most PafA-containing bacteria encode a singular *pafA* gene, some species encode two paralogous *pafA* genes (14), hinting that PafA could potentially form homo- or heterodimers in certain species.

Here we present biophysical and structural evidence that PafA_monomer_ and PafA_dimer_ coexist in solution. We make extensive use of hydrogen/deuterium exchange mass spectrometry (HDX-MS) to highlight the structural changes of PafA as it binds various nucleotides and dimerizes. Our data show that the N-terminal strand-helix motif and CTD become protected in the apo dimeric state, highlighting that these regions likely play a central role in maintaining the dimerization interface. We also leverage biochemical and structural prediction methods to demonstrate that the PafA_dimer_ populates a catalytically inactive state. These results provide critical insights into the structural dynamics and oligomeric heterogeneity of PafA.

## RESULTS

### PafA exists as a monomer and dimer in solution

During the purification of *Corynebacterium glutamicum* PafA (482 residues, 53.8 kDa) produced heterologously in *Escherichia coli*, we consistently observed two peaks corresponding to PafA in the final size-exclusion chromatography (SEC) step (**Fig. S3A** and **B**). We used SEC coupled to multi-angle light scattering (SEC-MALS) and identified two species with molecular weights corresponding to PafA_monomer_ (52 ± 1 kDa) and PafA_dimer_ (100 ± 5 kDa) (**Fig. S3C**). We employed analytical SEC to investigate the oligomerization behaviour of these PafA assemblies under different nucleotide-bound states. Our preparative-scale SEC column allowed us to isolate PafA_monomer_ and PafA_dimer_; however, approximately 10% of the other oligomeric forms remained as the column did not achieve baseline separation between the two peaks. We incubated isolated fractions of the preparative-scale PafA_monomer_ and PafA_dimer_ with or without nucleotide before conducting analytical SEC to investigate the interconversion of PafA across oligomeric states. These data reveal the PafA_monomer_ remained monomeric in the presence or absence of nucleotide (**Figure 1A** – top panels). By contrast, PafA_dimer_ (**Figure 1A** – bottom panels) is stabilized when bound to nucleotide but slowly dissociates in the apo state. The near-baseline separation of the two peaks in these analytical SEC experiments allowed us to readily deconvolute them and quantify the concentrations of PafA in the dimeric or monomeric states as a function of time. We fit these data to a kinetic model of monomer-dimer equilibrium (Equation 13 Supplemental Information) to obtain estimates of PafA dissociation (*k*_off_) and dimerization (*k*_on_) rates (**Figure 1B** and **C**). This analysis revealed that PafA_dimer_ dissociates into PafA_monomer_ with a *k*_off_ of 0.17 hour^-1^ (95% C. I. 0.02). While we obtained robust fits for *k*_off_, we could only obtain an upper bound of 1.6 ×10^4^ M^-1^ hour^-1^ for *k*_on_ (**Fig. S4**), corresponding to a lower bound for the dissociation constant (*K*_d_) of 1.1 mM.

**Figure 1.**
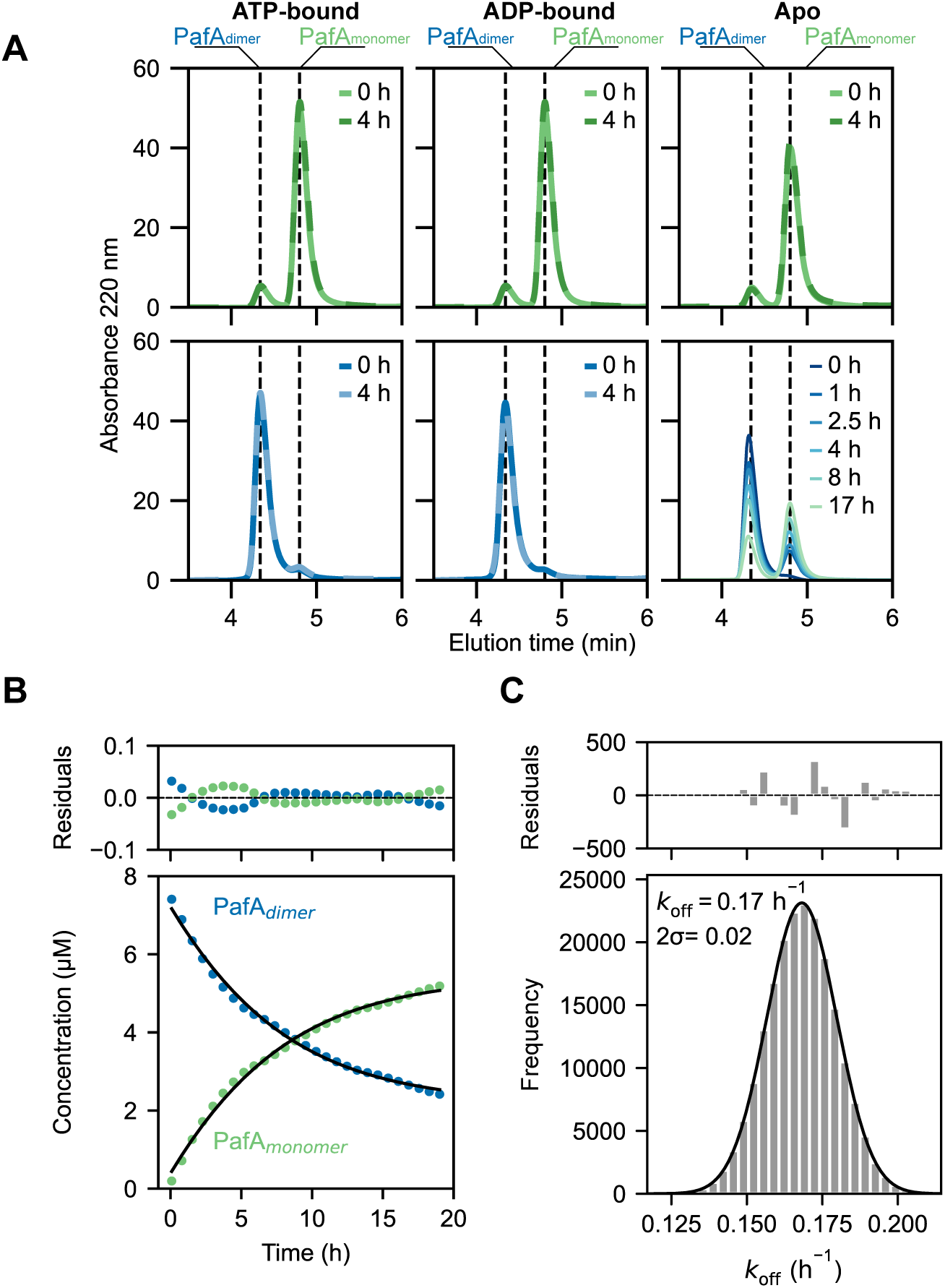
Investigating the oligomerization of PafA using SEC. **(A)** SEC traces of isolated PafA_monomer_ and PafA_dimer_ samples in the presence and absence of nucleotide. Left panels: ATP-bound PafA_monomer_ (green - top) and PafA_dimer_ (blue - bottom), middle panels: ADP-bound PafA_monomer_ (green - top) and PafA_dimer_ (blue - bottom), and right panels: apo PafA_monomer_ (green - top) and PafA_dimer_ (blue to green - bottom). PafA_monomer_ remains monomeric in the presence and absence of nucleotides. PafA_dimer_ is stabilized in the presence of nucleotide but dissociates in the apo state; **(B)** The concentrations of PafA_monomer_ (green) and PafA_dimer_ (blue) from the apo PafA_dimer_ sample (A – lower right panel) were plotted and the data were fit with an in-house derived dimerization - dissociation model (Supplemental Information); and **(C)** Histogram showing the *k*_off_ value and uncertainty extracted from 10,000 Monte Carlo simulations using the dimerization- dissociation model. The histogram (grey bars) displays the binned *k*_off_ values from this simulation. The histogram was fit to a Gaussian (black line). The center (*k*_off_) and 95% confidence interval (2 σ) were derived from the Gaussian fit.

### PafA_monomer_ and PafA_dimer_ have similar secondary structure but different activity

The slow interconversion rate of PafA_monomer_ and PafA_dimer_ allowed us to investigate their secondary structure and activity after SEC-based isolation. Circular dichroism (CD) spectra of PafA_monomer_ and PafA_dimer_ were indistinguishable, indicating no differences in secondary structure between these forms (**Figure 2A**). Next, we assessed the ability of PafA_monomer_ and PafA_dimer_ to pupylate the model target protein Log (4) by reconstituting the pupylation system *in vitro*, as previously described (25). We incubated isolated PafA_monomer_ and PafA_dimer_ with *C. glutamicum* Pup^E^, *Mycobacterium tuberculosis* Log, and ATP-Mg, and followed the pupylation reaction using sodium dodecyl-sulfate polyacrylamide gel electrophoresis (SDS-PAGE) (**Figure 2B**). Densitometry measurements of the pupylated Log (Log∼Pup; 41 kDa) protein band showed 3.6 ± 1.6 fold higher rate of Log∼Pup formation catalyzed by the PafA_monomer_ compared to PafA_dimer_ (**Figure 2C**).

**Figure 2.**
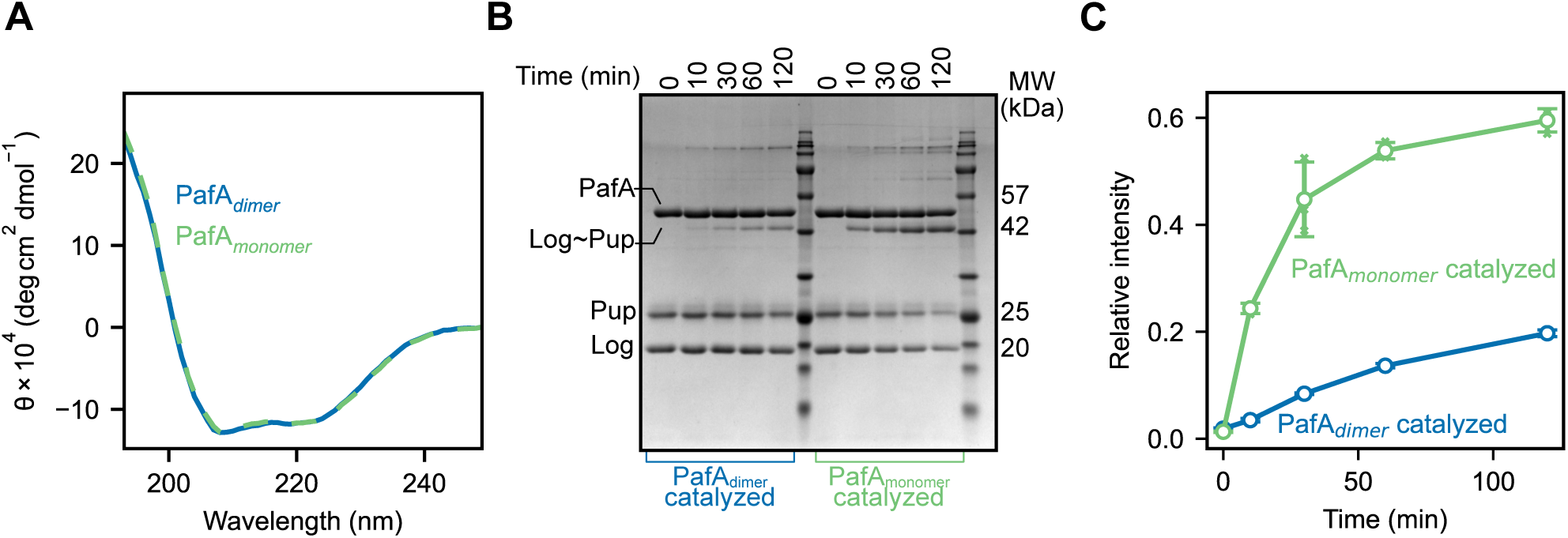
Biophysical and functional characterization of monomeric and dimeric PafA. **(A)** CD spectra of PafA_monomer_ (green) and PafA_dimer_ (blue); **(B)** SDS-PAGE displaying the pupylation of Log by PafA_dimer_ (left side - blue) and PafA_monomer_ (right - green); and **(C)** Densitometry analysis of the band corresponding to the molecular weight of pupylated Log from (B) was plotted. The band intensity of Log∼Pup in each reaction was measured relative to that of the corresponding PafA band at the 0-minute time point.

### Nucleotide binding induces global conformational changes in PafA

We used hydrogen-deuterium exchange mass spectrometry (HDX-MS) to investigate the conformational dynamics of PafA in various nucleotide-bound and oligomeric states. HDX-MS is a widely used technique for investigating protein structural changes and dynamics, particularly in the context of ligand binding and protein-protein interactions (26). HDX-MS monitors the exchange rate of amide hydrogens in protein backbone N-H groups with deuterium from the solvent, mediated by the transient opening and closing of hydrogen bonds (26, 27). Exchange incompetent ground-state amide groups (N-H_closed_) transiently visit exchange-competent higher-energy states (N-H_open_) and undergo H/D exchange according to:

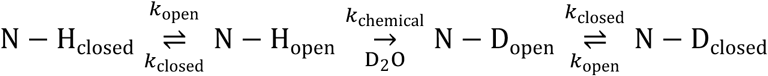

where *k*_open_ and *k*_closed_ are the rates of opening and closing events of individual amides, respectively, and *k*_chemical_ is the chemical or intrinsic H/D exchange rate of an unprotected amide. The observed H/D exchange rate, *k*_HDX_, for the reaction above is:

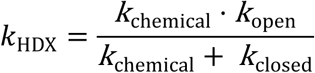

Hydrogen bonding is the primary factor that determines the HDX rate of a given amide. Protein segments that are heavily hydrogen bonded undergo slower HDX (up to 10^8^ fold) compared to dynamic regions, although access to solvent can also affect the HDX rate (28, 29). Allosteric transitions and protein/ligand binding can alter HDX rates by stabilizing or destabilizing specific regions of the protein. Binding events reduce HDX rates in the binding site, due to the formation of new H-bonds or decreased solvent accessibility. Allosteric effects can either increase or decrease HDX rates in distal regions of the protein by modulating protein dynamics, impacting the network of H-bonds or altering the solvent accessibility (30).

HDX follows two main kinetic regimes. In the commonly-observed EX2 regime (*k*_closed_ >> *k*_chemical_), HDX occurs after numerous opening and closing transitions. The overall HDX rate constant in the EX2 regime is:

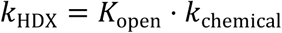

where

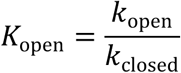

By contrast, in the EX1 regime (*k*_closed_ << *k*_chemical_), HDX occurs in the very first opening event and is indicative of concerted unfolding of amides, such that the HDX rate simplifies to the rate at which amides visit exchange-competent higher-energy conformations according to:

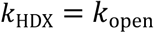

We began by using HDX-MS to investigate how nucleotide binding affects PafA_monomer_. We initially focused on continuous bottom-up labelling of the PafA_monomer_ in the apo-, ATP-bound, and ADP-bound states. Our peptide mapping of PafA yielded 189 peptides covering 97.5% of the protein sequence with a redundancy level of 4.7 (**Fig. S5**). The average H/D back exchange rate was 31% (**Table S1** (31) and **Fig. S6**). Most PafA peptides exhibited EX2 kinetics, with less than 15% displaying EX1 characteristics in some states. The peptides containing EX1 signatures will be extensively discussed in what follows.

For a global assessment of deuterium uptake patterns, we calculated the average deuterium uptake of peptides across all states and exposure times, then determined differential uptake by subtracting these values from one another and visualized the results using heat maps. Blue-coloured regions indicate a relative decrease in deuterium uptake while red regions show an increase. We colour-coded a previously published X-ray structure of PafA (15) to contextualize our HDX-MS results. ATP binding induced a striking decrease in deuterium uptake across virtually all structural elements of PafA_monomer_ (D_ATP- bound_ – D_apo_) (**Figure 3**). We observed the largest reduction in deuterium uptake in peptides encompassing residues that are directly involved in nucleotide binding or form the active site, spanning β1 (residues 13-22), β2 (residues 52-55), β3 (residues 58-63), β4 (residues 68-72), β6 (residues 128-136), β7 (residues 216-222), the loop between β3-β4 (residues 64-67), and loops in the CTD (413-417, 439-441 and 455-462). Interestingly, ATP binding also induced distal conformational changes in residues 260-295, 298-324, 331-343, and 354-478, up to 40 Å away from the nucleotide in the PafA_monomer_. These peptides encompass the peripheral helices of PafA and the entirety of the CTD, indicating that ATP binding has far-reaching effects on the conformation of the PafA_monomer_.

**Figure 3.**
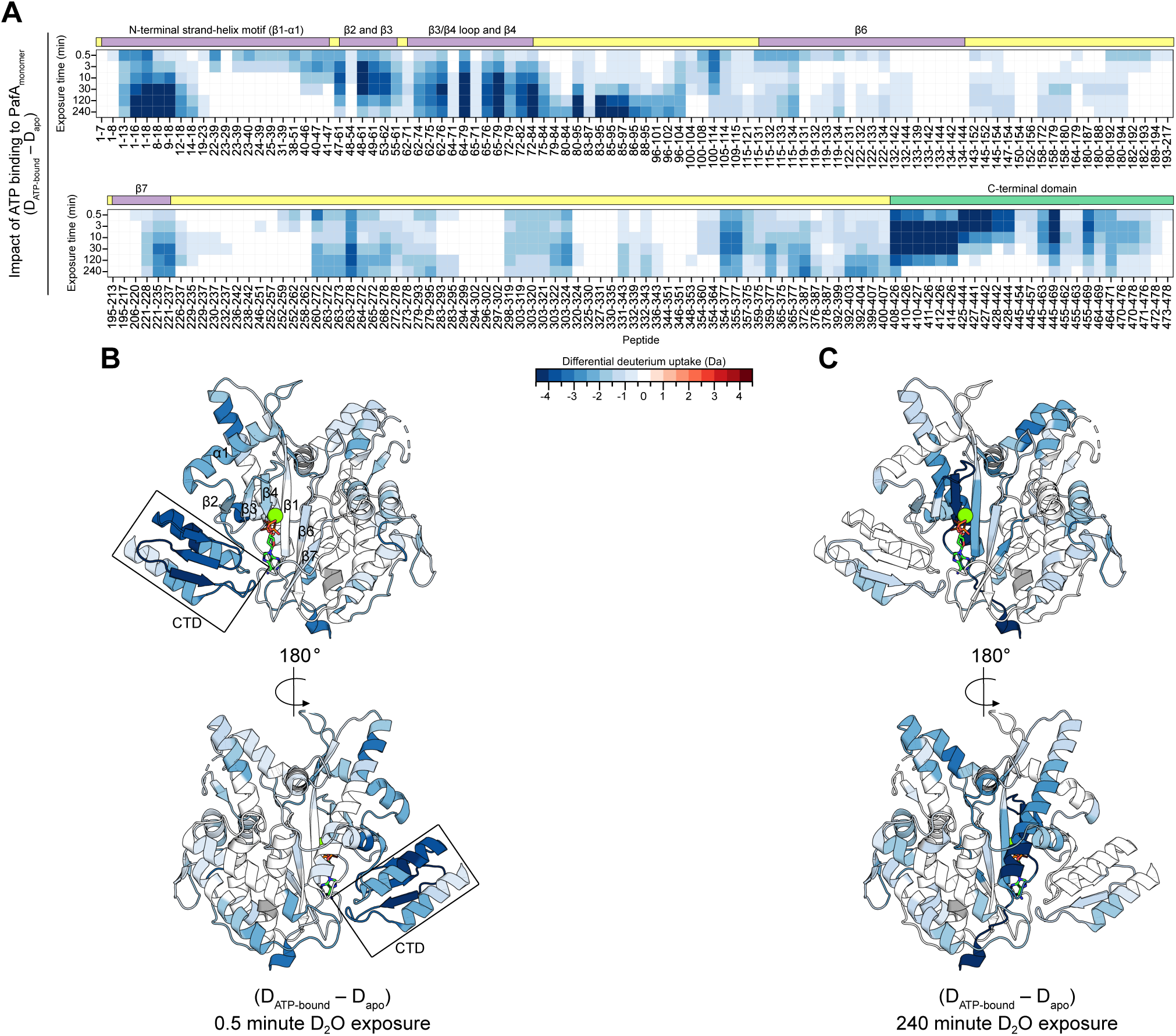
HDX-MS captures conformational changes in the PafA_monomer_ induced by ATP binding. **(A)** A heat map displaying differential deuterium uptake (D_ATP-bound_ – D_apo_) on the PafA_monomer_. Rows correspond to the D_2_O exposure time, columns correspond to the PafA peptide, and each square represents the differential deuterium uptake of the given peptide at the respective exposure time; **(B)** The 0.5-minute and **(C)** 240-minute D_2_O exposures are mapped onto the PafA crystal structures (PDB 4BJR (15)).

We next compared the deuterium uptake of the two nucleotide-bound states of PafA_monomer_ (D_ADP-bound_ – D_ATP-bound_) to investigate further how nucleotide influences PafA conformational dynamics. PafA_monomer_ experienced a subtle increase in deuterium uptake at early D_2_O exposure times (**Fig. S7A** and **B**) in peptides covering β1 and β6, and peptides encompassing W440 that interact with the nucleotide sugar moiety (6). α-helix 1 of the N-terminal strand-helix motif (residues 32-49), along with residues 143-154 and 260-272, 20 Å and 30 Å from the nucleotide-binding site, respectively, had a subtle increase in deuterium uptake. Later time points revealed a significant decrease in deuterium uptake in peptides that include R418, which contacts the adenine ring of bound nucleotides (6) and more subtle decreases in β2-4 and the loop preceding β7 (residues 205-215), which form a portion of the active site (**Fig. S7A** and **C**). We also observed an increase in deuterium uptake in distal peptides spanning residues 260-295, approximately 30 Å away from the nucleotide, at longer D_2_O exposure times. These data highlight conformational changes in PafA_monomer_ as it transitions between the ATP- and ADP-bound states, with a combination of local and global changes in deuterium uptake patterns.

### HDX-MS identifies regions involved in PafA dimerization

Given the effectiveness of HDX-MS in detecting changes in the conformational dynamics of PafA_monomer_ in response to ligand binding, we next applied this approach to examine the structural differences between the monomeric and dimeric forms of PafA in various nucleotide-bound states. This was possible because of the slow interconversion rate of PafA_monomer_ and PafA_dimer_ as indicated by our SEC-based experiments (**Figure 1**). As before, we performed bottom-up continuous labelling of the ATP- and ADP-bound states of PafA_dimer_ and PafA_monomer_ under the same conditions to enable comparative HDX-MS analysis. We observed minimal changes in deuterium uptake kinetics at early D_2_O exposure times between monomeric and dimeric PafA in the ATP-bound state (D_ATP-bound dimer_ - D_ATP-bound monomer_) (**Fig. S8A** and **B**). At longer exposure times, we observed moderate protection in parts of the active site (β2-4, and the β3-4 loop) and a portion of the CTD (residues 425-469), and high protection in CTD residues 408-426 (**Fig. S8A** and **C**). By contrast, residues 260-293 that encompass a peripheral α-helix had elevated deuterium uptake (i.e. more dynamic in PafA_dimer_ than in PafA_monomer_). Overall, our HDX data showed greater differences between monomeric and dimeric PafA in the ATP-bound state (D_ATP-bound dimer_ - D_ATP-bound monomer_) than in the ADP-bound state (D_ADP-bound dimer_ - D_ADP- bound monomer_), where we only noted a subtle decrease in deuterium uptake of CTD residues 425-444 at early D_2_O exposures (**Fig. S8A**, **D**, and **E**). Through this analysis, we identified several regions involved in PafA dimerization, particularly within the CTD and active site. However, since nucleotide binding significantly stabilizes both the monomer and the dimer, it might obscure structural differences between them in their nucleotide-bound states, as measured by HDX-MS.

To overcome the masking effects of nucleotide binding when examining the PafA dimerization interface, we compared apo PafA_monomer_ and apo PafA_dimer_ (D_Apo monomer_ - D_Apo dimer_). Our SEC data (**Figure 1**) helped guide the experimental design, revealing that apo PafA_dimer_ dissociates at *k*_off_ of 0.17 ± 0.02 h^-1^. Although the dissociation rate of apo PafA_dimer_ is slow, it is still sufficiently high to introduce structural heterogeneity via monomer formation during the course of our HDX experiment. Consequently, the deuterium uptake profile of apo PafA_dimer_ is likely to be contaminated by the monomer data, as the intact dimeric fraction diminishes in favour of an increasing proportion of dissociated monomers. To minimize this issue, we did not measure HDX kinetics over time. Instead, we only prepared a 30-second D_2_O exposure of apo PafA_dimer_ to assess the differential deuterium uptake of a sample representative of a true dimeric state with minimal dissociated PafA_monomer_ present compared to apo PafA_monomer_. Our HDX-MS data revealed more drastic differences in the apo state (D_Apo monomer_ - D_Apo dimer_) compared to the nucleotide-bound states (**Figure 4A** and **C**). Consistent with the nucleotide-bound HDX-MS results, we observed differential deuterium uptake in β2-4 of the active site and a significant increase in solvent-accessibility of CTD residues 408-444. We also observed a moderate increase in deuterium uptake in peptides spanning the N-terminal strand-helix motif (residues 1-51) and in CTD residues 445-478. These data highlight important structural elements that contribute to the PafA dimerization process.

**Figure 4.**
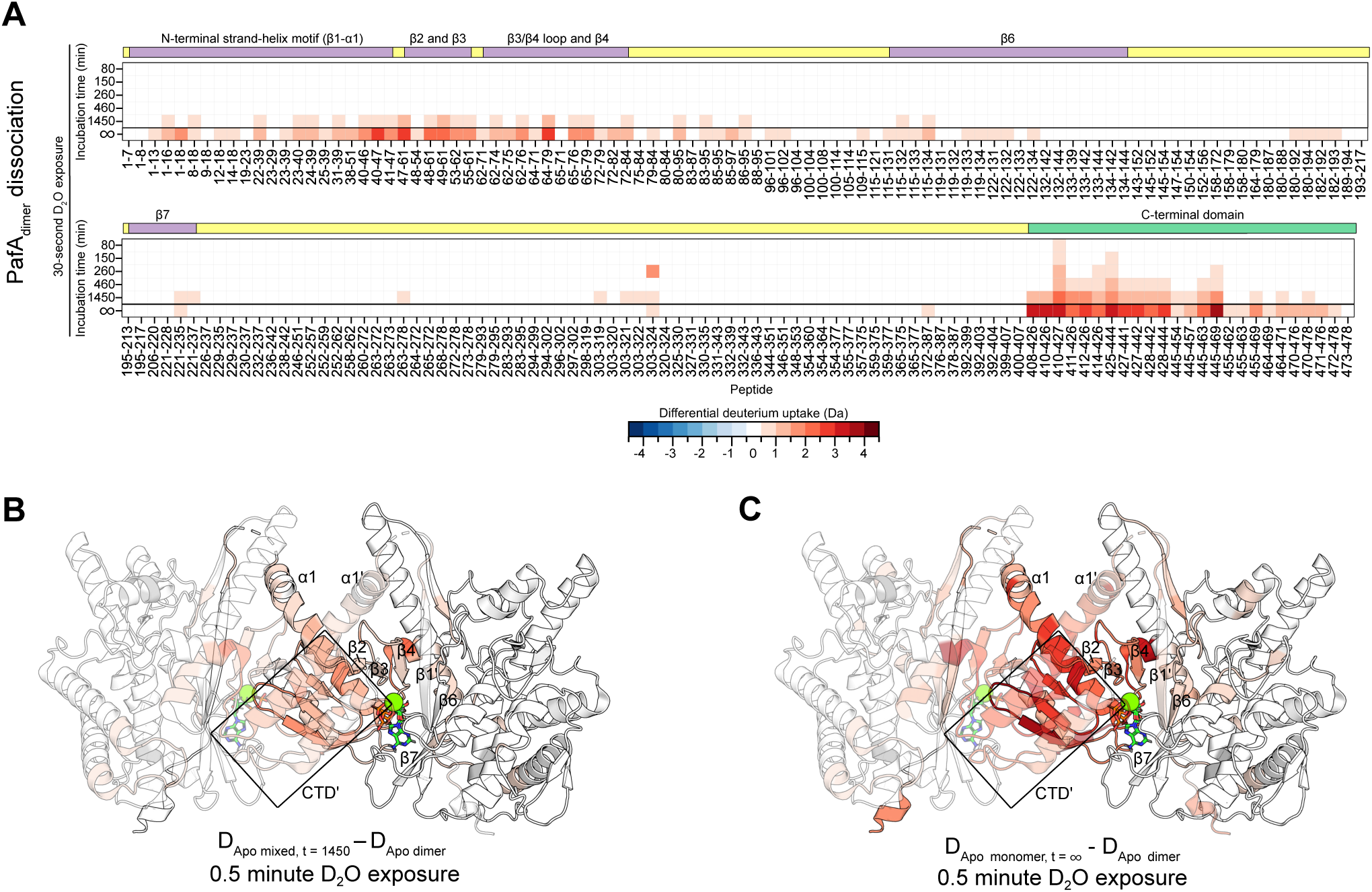
The CTD is involved in PafA dimerization. **(A)** A heat map displaying differential deuterium uptake between the apo PafA_monomer_ and PafA_dimer_ (D_Apo monomer_ - D_Apo dimer_) – denoted as t = ∞ on the bottom, and the dissociation of the apo PafA_dimer_ on the top (D_Apo mixed, t = 1450_ – D_Apo dimer_) of each segment. Rows correspond to the PafA peptide, columns correspond to the D_2_O exposure time, and each box represents the differential deuterium uptake of the given peptide at the respective exposure time. Differential deuterium uptake after 30 seconds of D_2_O exposure are mapped onto the PafA crystal structure (PDB 4B0T (6)); **(B)** PafA_dimer_ dissociation (D_Apo mixed, t = 1450_ – D_Apo dimer_) and; **(C)** The difference between the PafA_monomer_ and PafA_dimer_ (D_Apo monomer_ - D_Apo dimer_)highlight structural changes in the apo-state and identify the putative dimerization interface. One subunit in the structure is shown with 50% transparency to indicate the PafA_dimer_ dissociation process.

To further investigate the PafA_dimer_ dissociation process, we performed bottom-up pulsed HDX-MS experiments where we allowed PafA_dimer_ to dissociate for varying time periods before applying a 30-second D_2_O pulse to capture snapshots of how the relative abundance of PafA_dimer_ and PafA_monomer_ change over time. The D_Apo monomer_ - D_Apo dimer_ value measured above can be thought of as a *t* = ∞ endpoint of this process. Using the *k*_off_ value derived from our SEC data (**Figure 1**), we determined incubation time periods of 80, 150, 260, 460, and 1450 minutes to generate mixed PafA population containing >95, 85, 75, 65, and <10 percent of the apo PafA_dimer_ remaining in solution. When comparing the sample with the lowest PafA_dimer_ fraction (<10%) to the one with the highest (>95%) (D_Apo mixed, t = 1450_ – D_Apo dimer_), we noted an increase in deuterium uptake of the N-terminal strand-helix motif, β2-4, the loop between β3-4, and residues 408-471 of the CTD (**Figure 4A** and **B**). We also observed an increase in deuterium uptake in residues 303-324, located ∼10 Å from the CTD behind the active site cleft. These data are in close agreement with our findings on the structural differences between oligomeric forms in the apo state (D_Apo monomer_ - D_Apo dimer_) (**Figure 4A** and **C**), highlighting the regions that undergo a structural change as PafA transitions between the dimeric and monomeric state and further supporting the role of the CTD in this process.

### HDX-MS quantifies PafA dimer dissociation rate

We examined the raw mass spectra that were processed to generate our heat maps in the pulsed HDX experiments (**Figure 4A**). Most peptides showed symmetric isotopic distributions, indicative of EX2 kinetics. In contrast, some peptides in the N terminus and CTD displayed bimodal isotopic distributions, characteristic of slowly interconverting systems typical of EX1 kinetics (**Figure 5A**). Peptides with asymmetric spectra from the apo PafA_dimer_ state were chosen for in-depth analysis. These included peptides from the N-terminal domain encompassing the strand-helix motif and β2-3 (peptides 1-16, 1-18, 8- 18, and 9-18 covering β1, and 48-61, 49-61, and 55-61 spanning α1 and β2-3), and part of α2 less than 10 Å behind β1/3 (peptides 85-95 and 88-95). Notably, several peptides spanning the entire CTD (408-426, 410-426, 411-426, 412-426, 414-426, 427-442, 428-444, 445-463, 445-469, 470-476, 470-478, and 471-476) showed bimodal isotopic distributions, underscoring unique kinetic features in this region. We fit these bimodal isotopic distributions with Gaussian functions to compute the relative fraction of the protected and open populations (**Figure 5A**). These data were fit to a first-order exponential decay model to determine *k*_open_, thereby providing kinetic insights into the dynamic processes governing PafA_dimer_ dissociation in a site-specific manner (**Figure 5B**). The average *k*_open_ derived from our HDX data for PafA_dimer_ dissociation is 0.06 ± 0.06 h⁻¹, which is slightly lower than the *k*_off_ value derived from our SEC-based experiments (0.17 ± 0.02 h⁻¹ – **Figure 1**). The consistency in *k*_open_ values across these peptides , and their agreement with *k*_off_, suggests that the concerted unfolding of amides, characterized by the bimodal distributions, suggests that the concerted unfolding of amides, indicated by the bimodal distributions in the HDX mass spectra, reflects the same underlying process: PafA_dimer_ dissociation (**Figure 5B** and **C**). Importantly, these findings highlight the involvement of residues 1-18, 48-61, 85-95, and the CTD (residues 411-482) in PafA_dimer_ formation. We also observed bimodal distributions within the same peptides discussed above in the apo and ATP-bound PafA_monomer_ states (**Fig. S9)**. As earlier, we determined the *k*_off_ for these peptides. In the apo state, PafA_monomer_ can more readily sample higher energy states in the protein landscape, than in ATP-bound PafA_monomer_. Taken together, these results demonstrate that PafA samples similar high-energy intermediates in various ligation and oligomeric states and provide valuable insights into the previously uncharacterized dimerization interface of PafA in solution.

**Figure 5.**
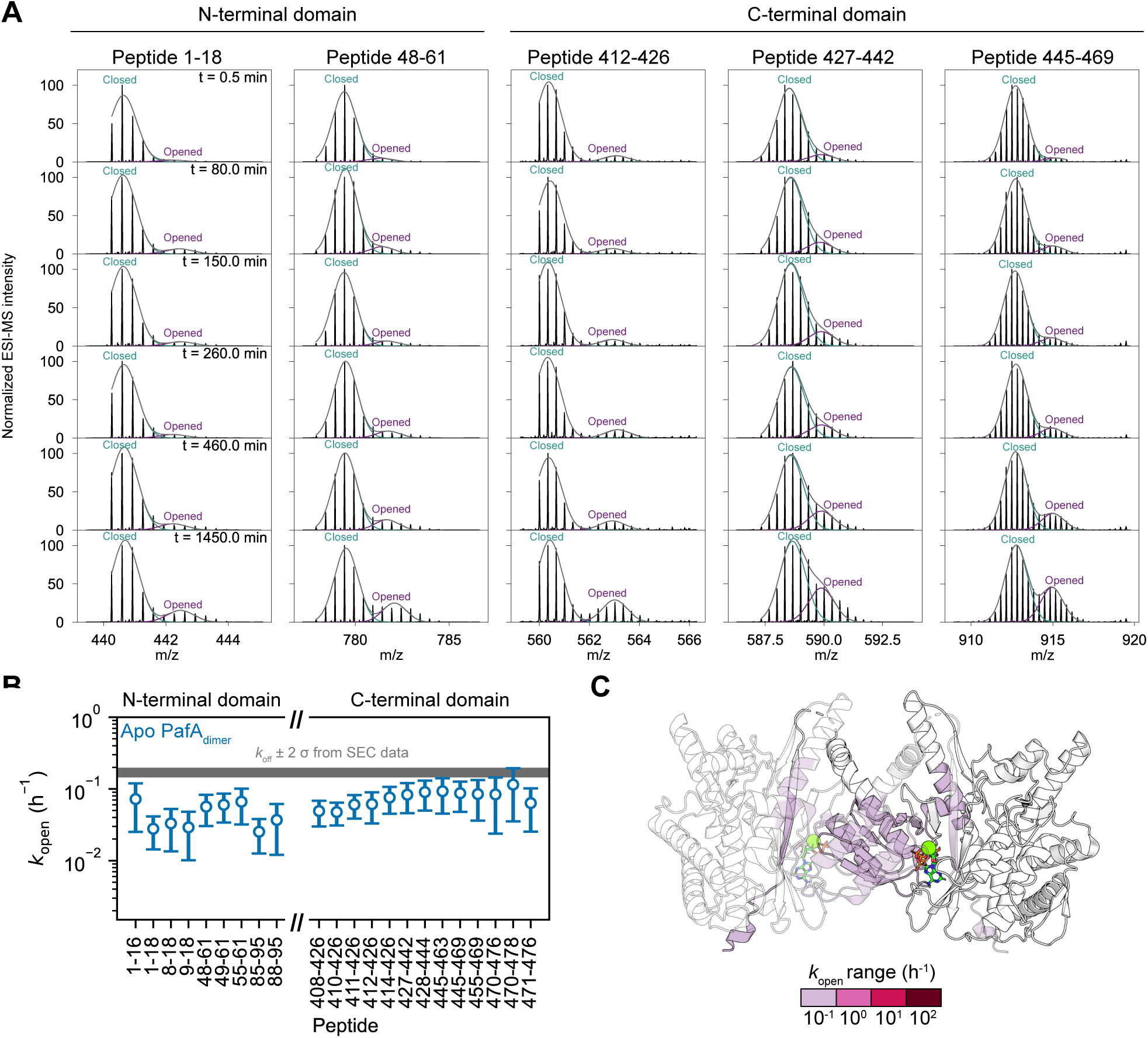
The PafA_dimer_ exhibits EX1 kinetics during the dimer dissociation process. **(A)** Representative isotopic spectra were fit with Gaussian distributions and the relative dimeric fraction was determined. The more protected distribution with a lower m/z represents the PafA_dimer_ and the less protected distribution with a higher m/z is representative of the dissociated PafA_monomer_; **(B)** The *k*_off_ values (h^-1^) per peptide are plotted. These represent the dissociation rate constants of regions involved in the PafA_dimer_ dissociation, with consistent values suggesting uniform structural transitions in these areas; **(C)** Peptides with asymmetric isotopic distributions are highlighted on the PafA crystal structure (PDB 4B0T (6)). One subunit is shown with 50% transparency to indicate the PafA_dimer_ dissociation process.

### AlphaFold3 does not predict a domain-swapped PafA_dimer_

We observed a slow timescale dissociation of PafA_dimer_ using SEC and HDX data, consistent with a domain-swapped dimeric structure (**Figure 1**, **4,** and **5**). Our HDX-MS data were interpreted in the context of the only available dimeric PafA structure, which features a domain swap involving the N-terminal strand-helix motif (β1/α1) to form the dimer(6, 32). To further explore the structural landscape of PafA_dimer_ subunit arrangements that are consistent with our HDX-MS data but are not represented in experimental structures, we used AlphaFold3 (AF3) computational modelling (33). We generated 6,000 structural models of PafA_dimer_ in various functional states, including apo, ATP-bound, ADP-bound, ATP-bound with Pup, ATP-bound with Log, and the quaternary complex with ATP, Pup and Log. We performed principal component analysis (PCA) on the AF3-generated models to examine structural variability among these. Clustering the dataset produced nine distinct clusters (**Fig. S10A**). We compared the exemplar structures of these clusters by aligning one PafA subunit from each dimeric structure (**Fig. S10**). The individual PafA subunits consistently adopted similar tertiary folds, with structural variations within 1 Å RMSD. However, each cluster exhibits a distinct subunit-subunit orientation. Notably, clusters A–C, which together comprise 65% of the dataset, form different dimerization interfaces that render PafA catalytically inactive (**Fig. S10B**, **C**, and **D**). Interestingly, none of the AF3-predicted structures exhibited a domain-swapped arrangement. Given the lack of an AF3-predicted domain-swapped structure, we believe that interpreting our HDX-MS data in the context of the only available X-ray structure of PafA_dimer_ is appropriate (**Figure 4** and **5**).

## DISCUSSION

The PPS is essential for the survival of pathogenic mycobacteria (1, 34). As such, members of the PPS are candidates for the development of new antibiotics. The lack of homology between PafA and functionally analogous proteins in the UPS makes PafA an attractive antibiotic target (6, 14). While the UPS employs numerous E3 ubiquitin ligases, each involved in ubiquitinating a narrow range of proteins, PafA is the sole Pup ligase responsible for pupylation of hundreds of target proteins in mycobacteria (11, 13, 35, 36). Despite extensive efforts to map the pupylome, no recruitment patterns have been identified, likely due to the lack of sequence and structural similarities among pupylation targets (36, 37).

Our HDX-MS data revealed that PafA is a highly dynamic enzyme that undergoes conformational fluctuations over a wide timescale in the apo and ATP-bound states. The conformational plasticity of PafA likely contributes to its function and ability to recruit diverse pupylation targets. We showed that recombinant PafA adopts both a monomeric and a dimeric conformation in solution. We demonstrate that PafA_monomer_ and PafA_dimer_ display different rates of pupylation *in vitro* against the model substrate Log. Therefore, mutations affecting PafA’s oligomeric state should be considered when assessing the functional role of residues, as changes in activity may result from altered oligomerization state rather than direct effects on ligand binding or target interactions. For example, recent work identified three electrostatic surface patches that facilitate target protein recognition (38). Our HDX-MS data show that the two patches in α1 (R32 and R38) and the CTD (R442 and K444) become protected in PafA upon dimerization. The dual role of these residues in substrate binding and PafA dimerization suggests an interplay between substrate recognition and the stabilization of the dimerization interface, potentially linking oligomeric state to the modulation of pupylation activity.

Our insights could be leveraged to advance structural studies. The two crystal structures of PafA bound to nucleotide and Pup [PDB 4B0T (6) and 4BJR (15)] have been pivotal in advancing our understanding of the structure: function relationship of this enzyme. However, important questions regarding the mechanisms of target protein selection and recruitment remain. This is compounded by the fact that no experimental structure of PafA bound to target proteins has been determined to date. In the domain- swapped PafA_dimer_ structure, the subunits associate and hinder the binding of the lower-affinity target proteins. Monomerization of PafA may facilitate target protein interactions and enable co-crystallization of PafA with its target proteins, advancing the understanding of how PafA interacts promiscuously with target proteins. Understanding the biological significance of these oligomeric states could provide valuable clues for developing small molecules that disrupt pupylation in pathogenic mycobacteria.

## EXPERIMENTAL PROCEDURES

### Plasmids and protein production

All genes were codon-optimized for expression in *Escherichia coli*. pET24a (Novagen, NJ) plasmids encoding for N-terminal His_6_-SUMO tagged *Corynebacterium glutamicum* PafA and Pup^E^ were obtained from Bio Basics Canada (Vaughn, ON) and a pET24a plasmid encoding for C-terminal His_6_-tagged *Mycobacterium tuberculosis* Log was obtained from Twist Bioscience (San Francisco, CA).

The plasmids were transformed into T7 Express *lysY* competent *E. coli* (New England Biolabs, MA). The cells were grown in LB broth (10 g/L NaCl, 10 g/L peptone, 5 g/L yeast extract), and the broth was supplemented with 100 mg/L MgCl_2_ for the growth of cells encoding the PafA construct. Cells were grown at 37 °C with 180 rpm agitation to an optical density (OD_600_) of 0.6-0.8 and were subsequently induced with 0.2 mM isopropyl-β-D- 1-thiogalactopyranoside (IPTG). Protein overproduction was allowed to proceed overnight (18-20 hours) at 30 °C for PafA or 18 °C for Log, and two hours at 37 °C for Pup. Following the induction period, all cells were harvested by centrifugation at 5000 × *g* relative centrifugal force (RCF) for 30 minutes at 4 °C. The cell pellets were resuspended in buffer A (50 mM Tris-HCl (pH 8.0), 300 mM KCl, 20 mM imidazole, and 10% [v/v] glycerol) and frozen at -20 °C.

### Protein purification

Resuspended cell pellets were thawed in lukewarm water. For cells containing His_6_- SUMO-PafA or His_6_-Log, cell lysis was achieved by homogenization in an EmulsiFlex-C3 (Avestin, CA) at 1,000 bar. For cells containing His_6_-SUMO-Pup, cells were lysed via boiling. The homogenate was clarified by centrifugation at 30,000 × *g* RCF for 30 minutes at 4 °C. The supernatant was loaded onto an immobilized metal affinity chromatography (IMAC) column with chelating sepharose beads (Cytiva, MA) charged with Ni^2+^. Column washes were performed with buffer A and buffer A supplemented with 50 mM imidazole. The protein was eluted with buffer A supplemented with 500 mM imidazole. His_6_-SUMO- PafA was incubated with Ubiquitin-like specific protease 1 (Ulp1) overnight at 4 °C, while being dialyzed into buffer W supplemented with 1 mM dithiothreitol (DTT). The cleaved protein was subjected to a second round of IMAC in which the flowthrough containing PafA was collected. All constructs were concentrated using centrifugal concentrators (Amicon, MO) with molecular weight cut-offs (MWCO) of 10 kDa for His_6_-SUMO-Pup and His_6_-Log, and 50 kDa for PafA. The concentrated proteins were loaded onto a Superdex 200 Increase 10/300 GL (Cytiva, MA) equilibrated in buffer B (50 mM Tris (pH 8.0), 150 mM NaCl, 20 mM MgCl_2_) (25). Fractions were collected and sample purity was evaluated via SDS-PAGE analysis. For His_6_-SUMO-Pup and His_6_-Log, fractions containing pure protein were pooled, mixed with 10% glycerol (v/v), flash-frozen in liquid nitrogen, and stored at -80 °C. The PafA_monomer_ and PafA_dimer_ fractions remained separate and were immediately used for subsequent experiments.

### SEC-MALS

PafA was buffer exchanged on a Superdex 200 Increase 10/300 GL column (Cytiva, MA) in 50 mM Tris (pH 7.5) and 100 mM NaCl. The PafA_monomer_ and PafA_dimer_ samples were pooled, filtered with a 2 µm filter, and prepared to 30 µM, and a 2 mg/mL BSA standard was run in tandem. An OMNISEC (Malvern Panalytical, Worcestershire, United Kingdom) system equipped with OMNISEC RESOLVE (CHR7100) Gel Permeation Chromatography (GPC) / Size Exclusion Chromatography (SEC), and OMNISEC REVEAL multi-detector modules (CHR6000) was utilized. Viscotex P 50 mm guard column and 300 mm analytical column were used in line with the OMNISEC system (Malvern Panalytical, Worcestershire, United Kingdom). Samples were stored at 7 °C in the autosampler and the columns were maintained at 20 °C. Data analysis was completed with the built-in OMNISEC software package.

### Analytical SEC

Separated PafA_monomer_ and PafA_dimer_ samples at 0.8 µM and 8.0 µM were prepared in Buffer B, with or without 5 mM of ADP or ATP supplemented. An Agilent 1100 series HPLC instrument (Santa Carla, CA) was equipped with a 150 mm ethylene bridged hybrid (BEH- 1.7 µm) 200 Å SEC column (Waters, MA). Buffer B also served as the running buffer with a flow rate of 0.3 mL/min. UV detection at 220 nm was recorded for the full duration of the 10-minute runs with a G1314A variable wavelength detector (Agilent, CA). Samples were injected with a G1367A autosampler (Agilent, CA) at ambient room temperature. The UV traces were exported from OpenLAB CDS LC ChemStation (Agilent, CA).

### SEC peak fitting

An exponentially modified Gaussian (EMG) model was used to analyze the chromatography peaks using equation 1 (39).

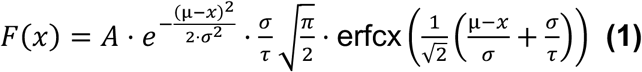

Here, *A* is the amplitude, µ is the center, σ is the standard deviation, and τ is the exponential parameter (39). Constraints were imposed for either the σ and τ values (analytical SEC) or the τ values (preparation scale SEC), setting them to be equal for the PafA_monomer_ and PafA_dimer_ peaks. The composite trapezoidal rule was utilized to determine the area under the two EMG curves using numpy.trapz. The goodness of fit was assessed by the reduced chi-square values.

### Deriving a two-state model for change in PafA oligomeric state

The reaction kinetics for a dimerization-dissociation process can be described by the following differential equation and its solution. The overall reaction is:

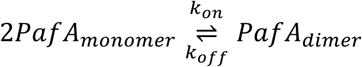

The concentration of each of the species at time : can be defined as:

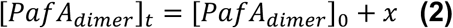

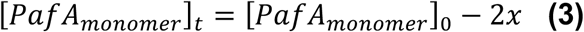

The rate equation can be written in terms of 1:

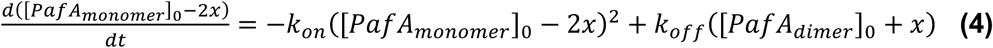

Separate the parameters and expand:

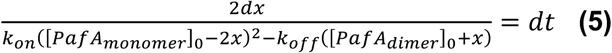

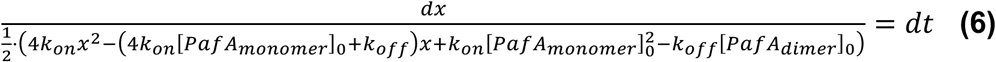

To simplify terms:

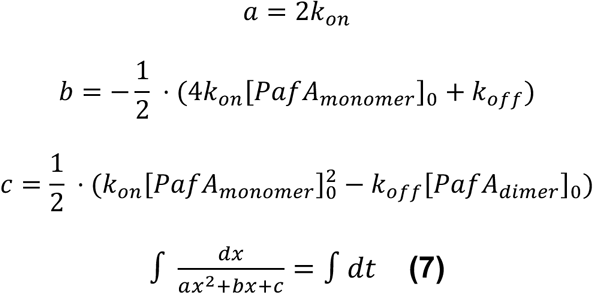

The solution to this differential equation is readily available from an integral table. To integrate the function 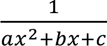 with respect to 1, we first complete the square in the denominator. The integral becomes:

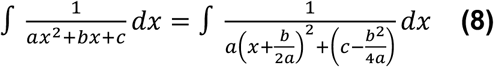

Depending on the discriminant Δ = D^7^ − 4=F, the integral can have different forms. Based on the biological constraints of this system, the discriminant must be greater than 0 (distinct real roots):

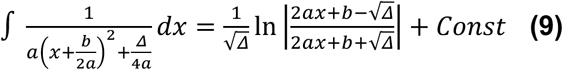

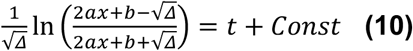

Here, *Const* is the constant of integration. This can be readily solved for using boundary conditions at : = 0, 1 = 0

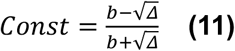

Therefore:

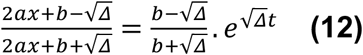

This can be rearranged to solve for *x*:

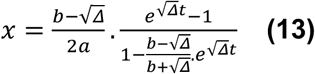

### Fitting change in [PafA_monomer_] and [PafA_dimer_] using the two-state model

The near baseline resolution of our analytical SEC enabled us to fit the relative fractions from the area under the EMG curves to the derived two-state dimerization-dissociation equation 13. To solve for [PafA_monomer_] and [PafA_dimer_] at a given time (*t*). For instances where NaN (not a number) was generated, that value was replaced with 0. Global fitting was performed using the 0.8 µM and 8.0 µM [PafA] datasets. A grid search was performed to explore combinations of the four fitting parameters: [PafA_monomer_]_0_, [PafA_dimer_]_0_, *k*_on_, and *k*_off_. Constraints were imposed to ensure *k_on_* and *k*_off_ values were equal between datasets in each grid search iteration. The goodness of fit was assessed by the reduced chi-square values.

### Monte Carlo simulations

Monte Carlo simulations were employed to determine confidence intervals for fitted parameters. Using the values from the best fit, ideal datasets were curated and residuals between the ideal dataset and raw input were calculated. The residuals were used to calculate the amount of random error, or noise, in the experimental data. The noise calculation was based on equation 14:

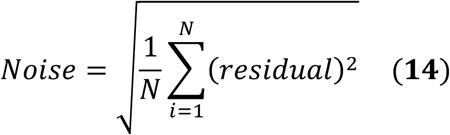

Where *N* is the number of data points. The *Noise* metric was used to introduce random perturbation from the ideal dataset to computationally mimic “experimental noise”. From here, the “noisy” data was subjected to the same fitting regime as the experimental dataset. This process was subsequently repeated 10,000-100,000 times.

### CD spectroscopy

PafA was separated using a Superdex 200 Increase 10/300 GL column (Cytiva, MA) in 25 mM Tris (pH 8.0) and 100 mM sodium phosphate. 3 µM isolated PafA_monomer_ and PafA_dimer_ samples were transferred into 1 mm pathlength cuvettes and circular dichroism spectrometry measurements were made with a JASCO J-815 circular dichroism (CD) spectropolarimeter (Tokyo, Japan). Measurements were made at 25 °C, scanning from 190-250 nm. Nine scans were summed, and an average spectrum was calculated.

### Pupylation assays

Following separation on a Superdex 200 Increase 10/300 GL column (Cytiva, MA), pupylation assays were prepared with isolated PafA_monomer_ and PafA_dimer_. 1000 µL reactions with 5 µM PafA, 20 µM His_6_-SUMO-Pup, and 10 µM His_6_-Log were carried out in Buffer B supplemented with 10 mM ATP at ambient room temperature (20-25 °C). Aliquots were quenched with an equal volume of Laemmli buffer and heat treatment. The samples were separated on a 15% sodium dodecyl-sulfate polyacrylamide gel electrophoresis (SDS-PAGE) gel. The SDS-PAGE gels were stained with Coomassie stain. Band density was analyzed using ImageJ (National Institutes of Health, Bethesda, MD, USA). Baseline optical density was removed using rolling ball background subtraction. Optical density plots for rows containing the substrate and product bands were obtained using the Gel Analyzer tool. Plots were manually divided into lanes and band densities were measured as the area under the curve for each lane. Results were normalized to the density of the PafA band and averaged across three replicate gels. The mean results fit an exponential growth model to determine the rates of the PafA_monomer_ and PafA_dimer_ catalyzed reactions, with error rates approximated by Monte Carlo.

### HDX-MS sample preparation

Continuous labelling bottom-up hydrogen-deuterium exchange mass spectrometry (HDX-MS) was performed on five states: (i) apo PafA_monomer_ (ii) ADP-bound PafA_monomer_, (iii) ATP-bound PafA_monomer_, (iv) ADP-bound PafA_dimer_, and (v) ATP-bound PafA_dimer_. Pulsed labelling bottom-up HDX-MS was performed on the apo PafA_dimer_. Buffer B was used as the H_2_O-based equilibrium buffer and D_2_O-based exchange buffer. The exchange buffer was adjusted to a pD 7.6 (pH_corr_ = 8.0) to account for the effect of D_2_O on glass electrode measurements (40, 41). During the equilibrium step, 8µM PafA_monomer_ or PafA_dimer_ were incubated with or without 5 mM of the respective nucleotide. HDX was initiated by a 10- fold dilution of the equilibrated stock into the exchange buffer, supplemented with 5 mM of nucleotide where appropriate. HDX was carried out in technical triplicate at room temperature (20-25 °C) with labelling times ranging from 0.5 to 1450 minutes for the continuously labelled samples, or 0.5 minutes per pulsed sample. HDX was quenched by acidification to pH_read_ 2.5 by mixing the sample 1:1 (v/v) with 250 mM NaH_2_PO_4_ (pH 2.23), 3 M guanidinium chloride, 3 mM n-dodecylphosphocholine (42), and 1.2% (v/v) formic acid, and flash freezing in liquid N_2_ followed by storage at -80 °C. A maximally deuterated sample was prepared as previously described (43), using PafA peptides acquired from online digestion with a nepenthesin-2 column (AffiPro, AP-PC-004, 1 mm × 20 mm) at 15 °C. Undeuterated samples were also prepared by mixing H_2_O-based sample with an equal volume of the quench buffer.

### HDX-MS liquid handling and mass analysis

Reverse-phase separation was performed on a M-class nanoAcquity UPLC instrument equipped with HDX technology (Waters, MA). 20 pmol of quenched sample was digested online with a nepenthesin-2 column (AffiPro, AP-PC-004, 1 mm × 20 mm) at 15 °C. The peptides were trapped on a BEH C18 (1.7 μm, 2.1 mm × 5 mm, Waters) column at 0 °C for 3 minutes with a flow rate of 100 µL/min. Following trapping, the peptides were separated at 0 °C on an HSS T3 column (1.8 μm, 1.0 × 50 mm, Waters) using an 8-minute linear acetonitrile: H_2_O gradient acidified with 0.1% formic acid, at a flowrate of 100 μL/min. The syringe and sample loop were cleaned between each injection to reduce carryover, using 1.5 M guanidine hydrochloride, 4% (v/v) acetonitrile, 0.8% (v/v) formic acid, and 1.5 mM n-dodecylphosphocholine.

The UPLC outflow was directed to a quadrupole ion mobility time-of-flight (Q-TOF) Synapt G2-Si mass spectrometer (Waters, MA), equipped with a standard electrospray source operating in positive-ion mode and a capillary voltage set to +3 kV. Instrument calibration was dynamically performed using LeuEnk solution (1+ ion, 556.2771 m/z), which was infused through the LockSpray capillary at a flow rate of 10 μL/min and sampled every 20 seconds. Drift time-aligned MS^E^ data-independent acquisition was utilized for peptide mapping, following established methods (44) . Data acquisition covered the *m/z* range of 50–2000, with a scan duration of 0.4 seconds per cycle. Fragmentation was achieved by varying the transfer collision energy linearly between 20 and 40 V during alternating scans. Ion mobility separation was manually controlled as outlined previously (45),and the quadrupole was tuned to isolate ions with *m/z* values above 300, excluding smaller ions from entering the mobility cell. The TOF mass analyzer was operated in resolution mode.

### HDX-MS data processing

The undeuterated MS^E^ data were analyzed with ProteinLyxn Global Server (version 3.0.3) (Waters, MA) to identify the PafA peptides present. Peptides were filtered in DyanmX (version 3.0) (Water, MA) using previously described filtering parameters (46), before HDX-MS data analysis. Peptides were manually inspected to ensure only high-quality spectra were retained for analysis. An in-house Python script was used to perform a hybrid significance test on the retained peptides and generate heat maps for the relative deuterium uptake between two states. The maximally deuterated samples were used to determine back-exchange rates during the liquid handling and mass analysis process (43).

### AlphaFold 3 modeling and categorization

AlphaFold 3(33) was used to predict the structures of PafA in six distinct states: (i) apo, (ii) ADP-bound, (iii) ATP-bound, (iv) ATP-bound with Pup, (v) ATP-bound with Log, and (vi) ATP-bound with both Pup and Log. Predictions involving nucleotide-bound states also included Mg ions. Each prediction was randomly seeded and yielded five models.

The alpha-carbon coordinates were extracted from the AlphaFold 3 models for PafA residues which displayed asymmetric isotopic distributions in the HDX-MS data (residues 1-18, 48,61, 85-95, and 408-478). All other chains or ligands were ignored. Principal component analysis was performed on the extracted coordinates using the ProDy software package (47). The dataset was projected onto the first three principal components, which together accounted for 82% of the total variance. The DBSCAN (Density-Based Spatial Clustering of Applications with Noise) algorithm from Scikit-learn (48) was then applied to identify and remove outliers, with the distance threshold (ɛ) set to 2 and the minimum number of points required to form a cluster set to 10. Following denoising, the first two principal components were retained and scaled according to their fraction variance explained (0.49 and 0.28 respectively). Affinity propagation clustering was performed on the scaled coordinates using the Scikit-learn library (48) to identify distinct modes of dimerization and obtain exemplar structures from each cluster. A maximum number of 1000 iterations was allowed to enable the model to converge. The preference was set to -200 for all points to encourage the formation of fewer clusters.

## DATA AVAILABILITY

Mass spectrometry data are available from the MassIVE database under entry MSV000096530.

## SUPPORTING INFORMATION

This article contains supporting information.

## ACKNOWLEDGEMENTS

A.P. acknowledges support from a Natural Sciences and Engineering Research Council of Canada Postgraduate Doctoral Scholarship. J.C.O.B. acknowledges support from a Queen Elizabeth II Graduate Scholarship in Science and Technology. Financial support was provided by a Canadian Institutes of Health Research Project Grant PJT451412 (to S.V.) and a Natural Sciences and Engineering Research Council of Canada Discovery Grant RGPIN-2021-02843 (to S.V.). MS data were recorded at the Mass Spectrometry Facility of the Advanced Analysis Centre, University of Guelph. We thank Dr. Dyanne Brewer (University of Guelph) for assistance with MS measurements. We thank Dr. Algirdas Velyvis (University of Guelph) for helpful discussions and critical review of the manuscript.

## AUTHOR CONTRIBUTIONS

A.P. and S.V. conceptualized the project. A.P. performed biochemical and biophysical experiments, and completed the AlphaFold3 structural predictions. J.C.O.B. performed densitometry measurements and bioinformatic analyses. A.P. and S.V. analyzed and interpreted data, and wrote the manuscript. S.V. supervised the research and secured funding for the study.

## ABBREVIATIONS

ADP: adenosine diphosphate
AF3: AlphaFold3
ATP: adenosine triphosphate
BEH: ethylene bridged hybrid
BSA: bovine serum albumin
C.I.: confidence interval
CD: circular dichroism
CTD: C-terminal domain
DBSCAN: Density-Based Spatial Clustering of Applications with Noise
DTT: dithiothreitol
HDX: hydrogen deuterium exchange
HPLC: high-performance liquid chromatography
IMAC: immobilized metal affinity chromatography
IPTG: isopropyl-β-D-1-thiogalactopyranoside
*K*_d_: equilibrium dissociation constant
*k*_off_: dissociation rate constant
*k*_on_: association rate constant
m/z: mass to charge ratio
MALS: multiangle light scattering
MS: mass spectrometry
MWCO: molecular weight cut-off
NTD: N-terminal domain
OD_600_: optical density
PCA: Principal Component Analysis
PDB: Protein Data Bank
PPS: Pup-proteasome system
Q-TOF: quadrupole-time of flight
RCF: relative centrifugal force
RMSD: root mean square deviation
RPM: rotations per minute
SDS-PAGE: sodium dodecyl sulphate–polyacrylamide gel electrophoresis
SEC: size exclusion chromatography
SUMO: small ubiquitin-like modifier
UPLC: ultra-performance liquid chromatography
UPS: ubiquitin-proteasome system
UV: ultraviolet

