## Supporting Information for "Probing the structural heterogeneity of Pup ligase PafA using H/D exchange mass spectrometry"

**This file contains the following materials:**

Supplemental Figure 1

Supplemental Figure 2

Supplemental Figure 3

Supplemental Figure 4

Supplemental Figure 5

Supplemental Figure 6

Supplemental Figure 7

Supplemental Figure 8

Supplemental Figure 9

Supplemental Figure 10

Supplemental Table 1

Supplemental References

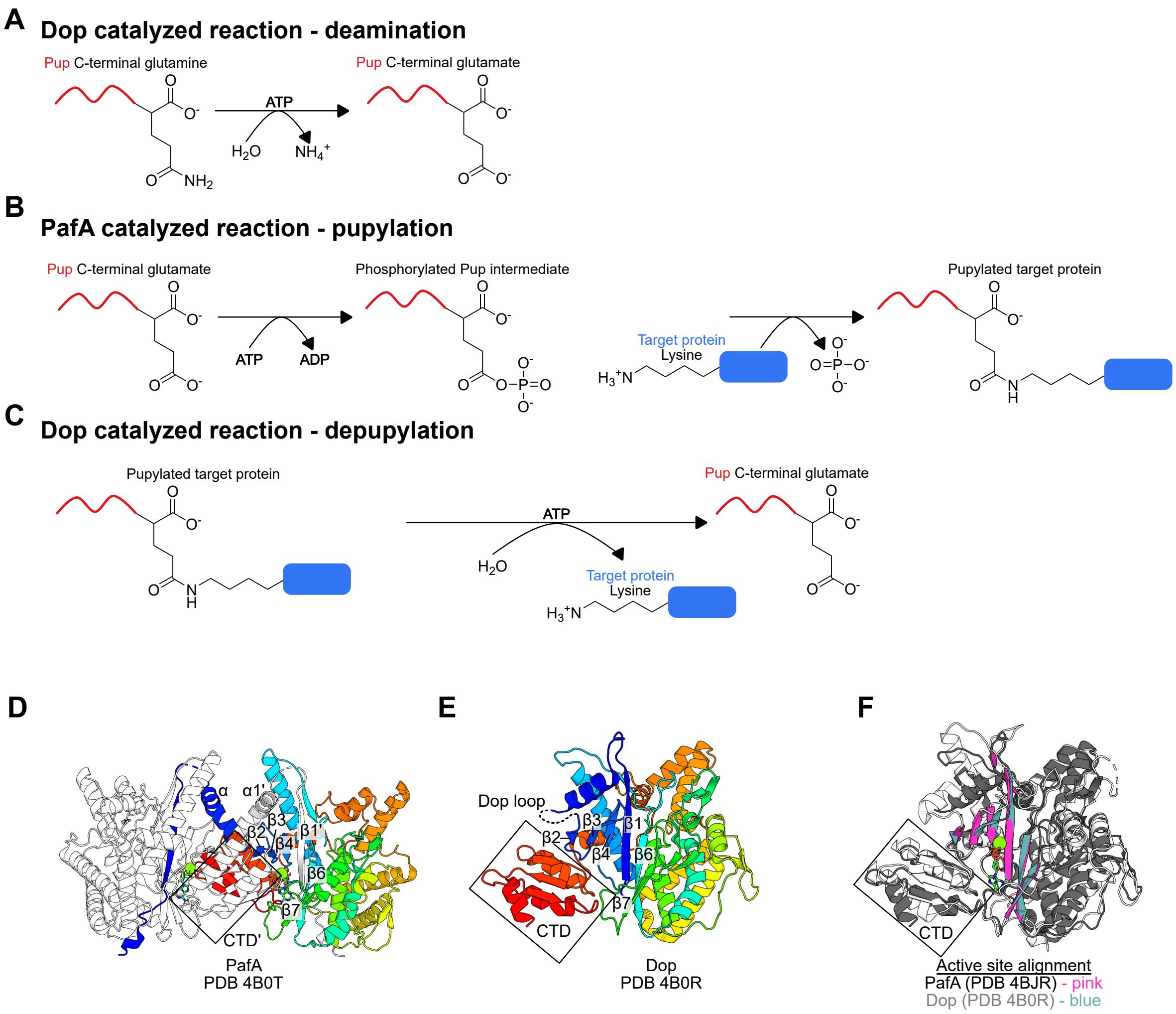


**Supplemental Figure 1. Comparison between PafA and Dop.** Schematics of **(A)** Dop catalyzed deamidation of Pup; **(B)** PafA catalyzed pupylation and; **(C)** Dop catalyzed depupylation. Dop retains ADP and inorganic phosphate in the active site during catalysis, resulting in sub-stoichiometric consumption of ATP (1). Whereas PafA consumes ATP stoichiometrically for pupylation (2); **(D)** Crystal structure the domain-swapped *Corynebacterium glutamicum* $\text{PafA}_{\text{dimer}}$ [PDB 4B0T (2)]. One subunit is coloured by chainbow, with the N-terminus in dark blue and C-terminus in red. The other subunit is white. The N-terminal strand-helix motif (β1/α1) undergoes a domain swap; **(E)** Crystal structure of Dop from *Acidothermus cellulolyticus* [PDB 4B0R (2)] coloured by chainbow, with the N-terminus in dark blue and C-terminus in red. The location of the disordered Dop loop is indicated; **(F)** Alignment of the *C. glutamicum* $\text{PafA}_{\text{monomer}}$ (white with pink active site β-sheet cradle) [PDB 4BJR (3)], and Dop [PDB 4B0R (2)] (grey with light blue active site β-sheet cradle).

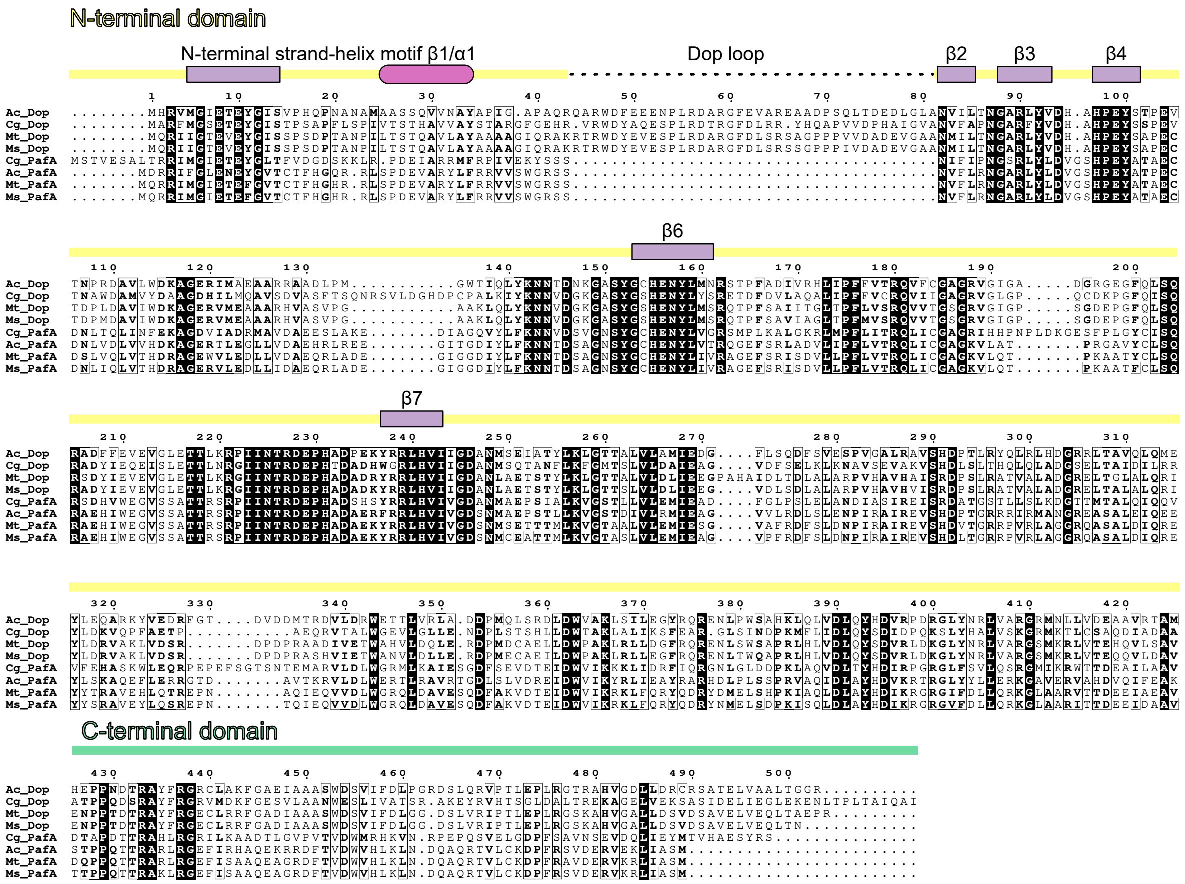


**Supplemental Figure 2. Sequence alignment of Dop and PafA**. A multiple sequence alignment was performed on a curated set of 80 Dop and PafA sequences from the UniProtKB database. The alignment of sequences for Dop and PafA from *Mycobacterium tuberculosis* (Mt), *M. smegmatis* (Ms), *Corynebacterium glutamicum* (Cg), and *Acidothermus cellulolyticus* (Ac) are displayed. Key structural features including the N-terminal strand-helix motif, the active site β-sheet cradle, Dop loop, and C-terminal domain are indicated (2).

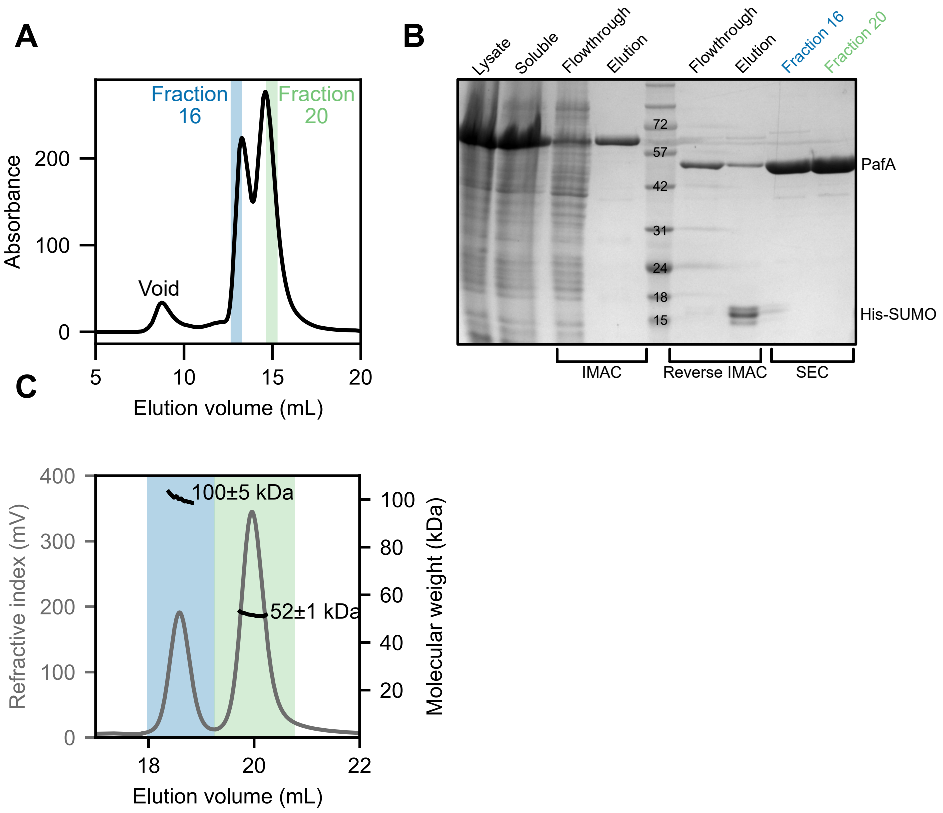


**Supplemental Figure 3. PafA coexists in monomeric and dimeric states during purification. (A)** Two distinct peaks are observed during the final size-exclusion chromatography step in PafA purification; **(B)** SDS-PAGE reveals the predominant protein in fractions 16 (blue) and 20 (green) in the two SEC peaks corresponds to the approximate molecular weight of PafA. These fractions are retained as $PafA_{dimer}$ and $PafA_{monomer}$ for downstream experiments, respectively; **(C)** SEC-MALS identified PafA monomers ($PafA_{monomer}$ - green) and dimers ($PafA_{dimer}$ - blue) in solution. The refractive index (grey) trace displays two distinct peaks with molecular weight (black) assignments corresponding to $PafA_{monomer}$ (54 kDa) and $PafA_{dimer}$ (108 kDa).

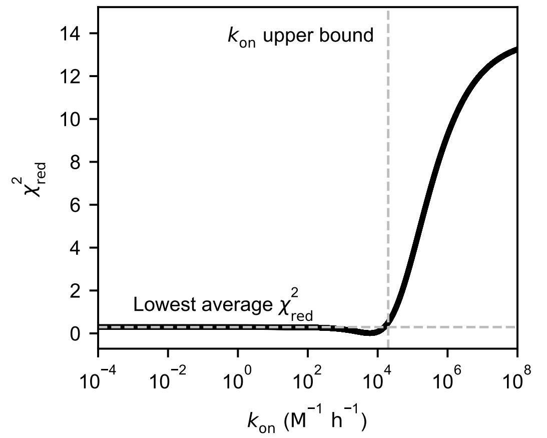


**Supplemental Figure 4. Determination of $k_{on}$ upper bound.** The best-fit parameters from the two-state dimerization-dissociation model for the two initial protein concentrations ($[PafA_{monomer}]_0$ and $[PafA_{dimer}]_0$) and $k_{off}$ were used to determine the upper bound for $k_{on}$. The reduced chi-squared ($X^2_{red}$) values were plotted against 10,000 $k_{on}$ fitted values. Reduced chi-square is insensitive to $k_{on}$ values ranging from 1 $\times 10^{-4}$ to 1.6 $\times 10^4$ $M^{-1}$ $hour^{-1}$ with a shallow minimum at 6 $\times 10^3$ $M^{-1}$ $hour^{-1}$.

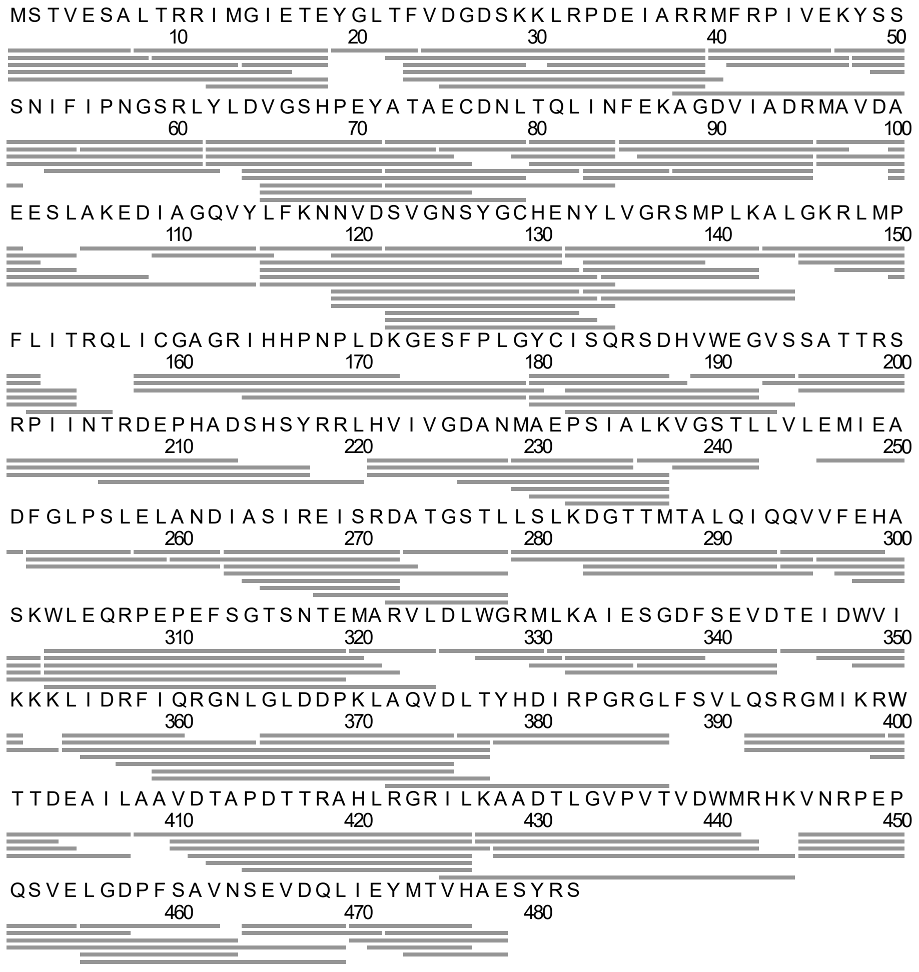


**Supplemental Figure 5**. Peptide coverage map for PafA. PafA peptide mapping yielded 189 peptides (grey bars) covering 97.5% of the sequence with a redundancy level of 4.7.

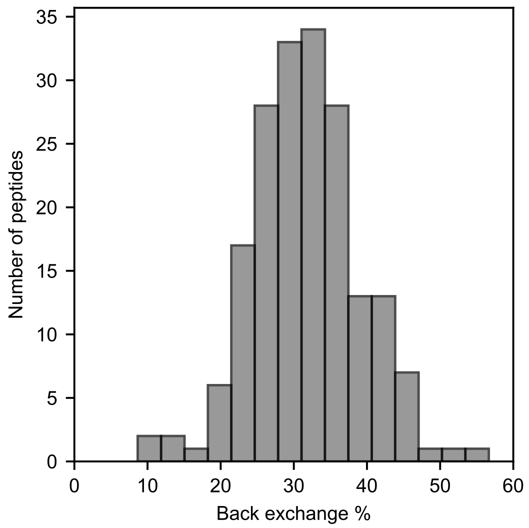


**Supplemental Figure 6. Histogram displaying the percentage of HDX back exchange for PafA peptides.**

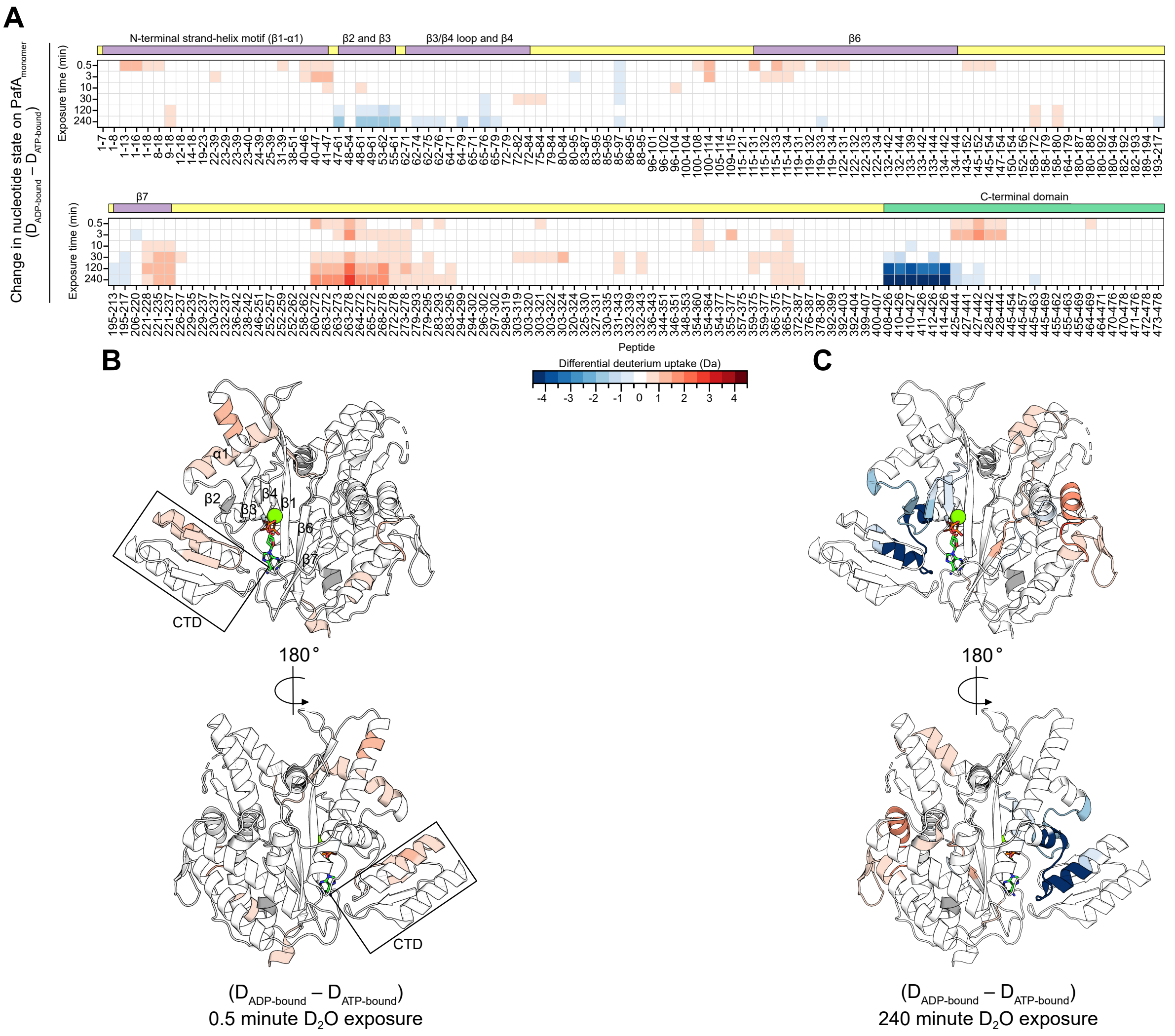


**Supplemental Figure 7. Change in nucleotide state impacts the conformational dynamics of the PafA$_{monomer}$. (A)** A heat map displaying differential deuterium uptake ($D_{ADP\text{-}bound}$ – $D_{ATP\text{-}bound}$) on the PafA$_{monomer}$. Rows correspond to the $D_2O$ exposure time, columns correspond to the PafA peptide, and each square represents the differential deuterium uptake of the given peptide at the respective exposure time; **(B)** The 0.5-minute and **(C)** 240-minute $D_2O$ exposures are mapped onto the PafA crystal structures [PDB 4BJR (3)].

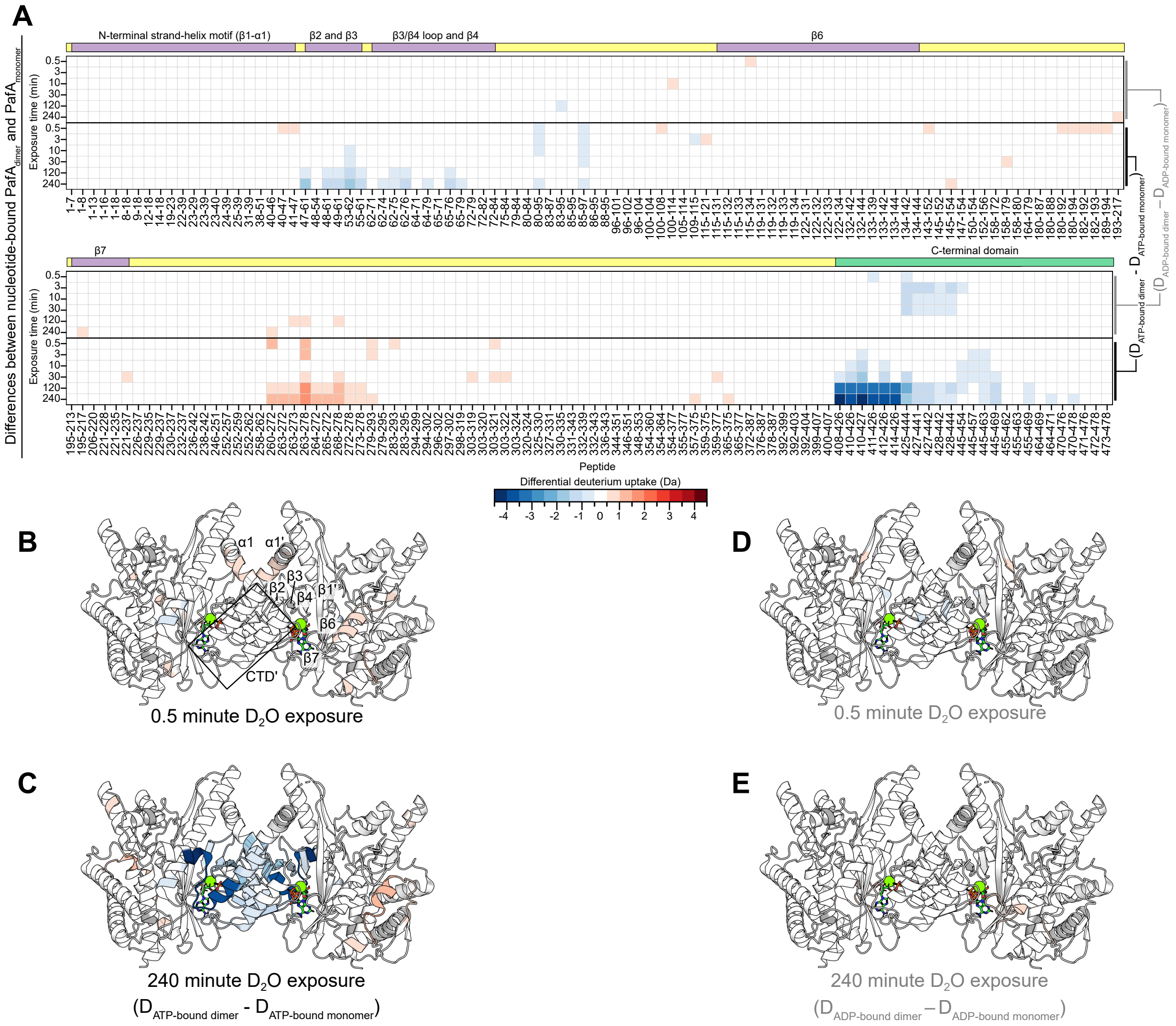

**Supplemental Figure 8. Differences in the nucleotide-bound PafA$_{dimer}$ and PafA$_{monomer}$ identify regions involved in dimerization. (A)** Heat maps displaying differential deuterium uptake of the nucleotide-bound PafA$_{dimer}$ ($D_{ADP\text{-bound dimer}}$ - $D_{ADP\text{-bound monomer}}$) on the top, and ($D_{ATP\text{-bound dimer}}$ - $D_{ATP\text{-bound monomer}}$) on the bottom of each segment, in reference to the PafA$_{monomer}$. Rows correspond to the $D_2O$ exposure time, columns correspond to the PafA peptide, and each square represents the differential deuterium uptake of the given peptide at the respective exposure time; **(B and C)** The 0.5-minute and 240-minute $D_2O$ exposures for $D_{ATP\text{-bound dimer}}$ - $D_{ATP\text{-bound monomer}}$ and **(D and E)** the 0.5-minute and 240-minute $D_2O$ exposures for $D_{ADP\text{-bound dimer}}$ - $D_{ADP\text{-bound monomer}}$ are mapped onto the PafA crystal structures [PDB 4B0T (2)].

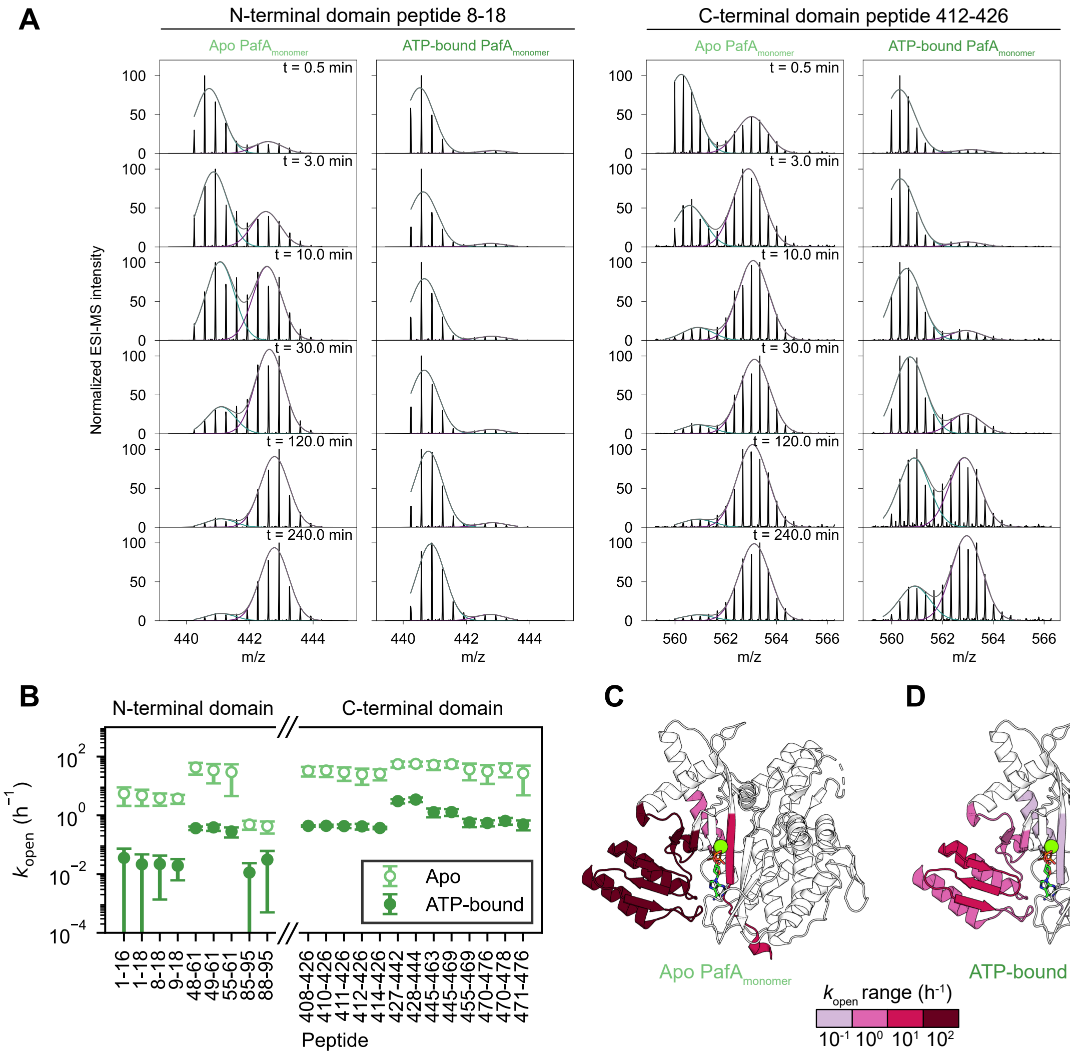


**Supplemental Figure 9. PafA undergoes cooperative unfolding events in the apo state. (A)** Representative peptides with asymmetric isoptic distributions for the apo and ATP-bound $PafA_{monomer}$ states were fit with Gaussian functions. The shift in the protected state with lower m/z to the open populations illustrates the kinetic differences between conformational states; **(B)** The $k_{off}$ values determined from the fits are plotted for each peptide. Distinct off-rates indicate variations in conformational stability, with ATP binding resulting in decreased $k_{off}$ values, compared to the apo $PafA_{monomer}$; **(C)** The $k_{off}$ values are colour-mapped onto the PafA structure [PDB 4BJR (3)].

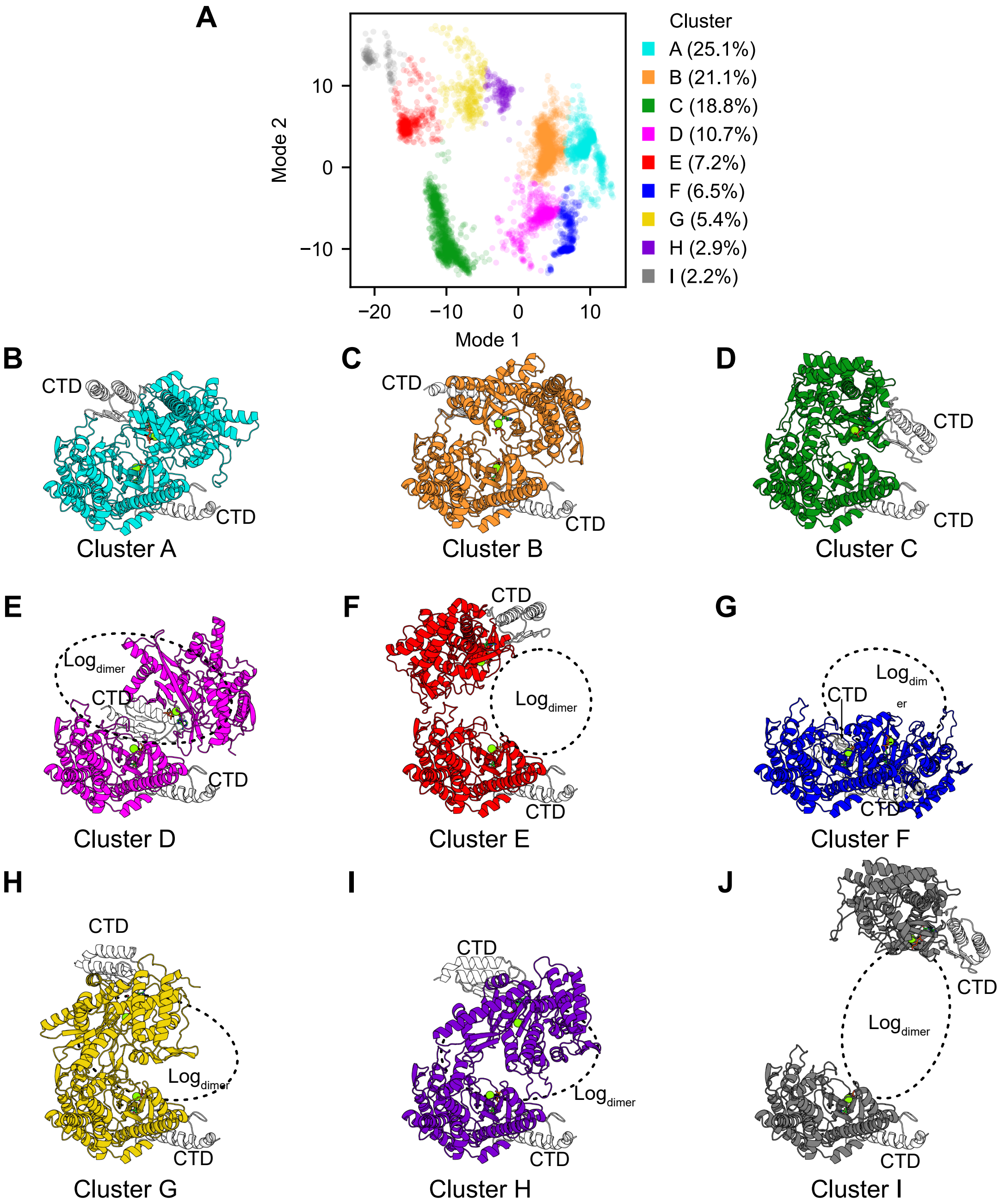


**Supplemental Figure 10. $PafA_{dimer}$ dimerization interfaces predicted by AlphaFold 3. (A)** Two-dimensional principal component plot. Clusters were determined with Affinity Propagation and are colour-coated;**(B-J)** The exemplar $PafA_{dimer}$ from Cluster A through I, respectively, are displayed. The N-terminal domain is coloured based on the cluster assignment in the 2D plot,and the CTD is shown in white to aid in identification. One subunit was aligned for all cluster exemplars. The area where Log resides is shown with a dotted circle.

### Supplemental Table 1. HDX summary

| | Apo $PafA_{monomer}$ | ATP-bound $PafA_{monomer}$ | ADP-bound $PafA_{monomer}$ | ATP-bound $PafA_{dimer}$ | ADP-bound $PafA_{dimer}$ | Apo $PafA_{dimer}$ |
|---|---|---|---|---|---|---|
| HDX reaction details | Final $D_2O$ concentration (v/v): 90 %<br>$pH_{corr}$ 7.6, room temperature | Final $D_2O$ concentration (v/v): 90 %<br>$pH_{corr}$ 7.6, room temperature, 5 mM ATP | Final $D_2O$ concentration (v/v): 90 %<br>$pH_{corr}$ 7.6, room temperature, 5 mM ADP | Final $D_2O$ concentration (v/v): 90 %<br>$pH_{corr}$ 7.6, room temperature , 5 mM ATP | Final $D_2O$ concentration (v/v): 90 %<br>$pH_{corr}$ 7.6, room temperature, 5 mM ADP | Final $D_2O$ concentration (v/v): 90 %<br>$pH_{corr}$ 7.6, room temperature |
| Pre-$D_2O$ exposure incubation time (min) | 10 | | | | | 0.5, 80, 150, 260, 460, 1450 |
| HDX time course (min) | 0.5, 3, 10, 30, 120, 240 | | | | | 0.5 |
| Undeuterated controls | 3 | | | | | |
| Back-exchange | 31 % (range: 8.6 – 56.7 %) | | | | | |
| Number of peptides | 189 | | | | | |
| Average peptide length/ redundancy | 10.7 ± 4.9 / 4.7 | | | | | |
| Replicates | 3 technical replicates | | | | | |
| Repeatability | 0.03 Da | | | | | |
| Significant difference | 0.5 Da | | | | | |

### SUPPORTING INFORMATION REFERENCES


1. Bolten, M., Vahlensieck, C., Lipp, C., Leibundgut, M., Ban, N., and Weber-Ban, E. (2017) Depupylase Dop Requires Inorganic Phosphate in the Active Site for Catalysis. *Journal of Biological Chemistry*. **292**, 4044–4053
2. Özcelik, D., Barandun, J., Schmitz, N., Sutter, M., Guth, E., Damberger, F. F., Allain, F. H.-T., Ban, N., and Weber-Ban, E. (2012) Structures of Pup ligase PafA and depupylase Dop from the prokaryotic ubiquitin-like modification pathway. *Nat Commun*. **3**, 1014
3. Barandun, J., Delley, C. L., Ban, N., and Weber-Ban, E. (2013) Crystal Structure of the Complex between Prokaryotic Ubiquitin-like Protein and Its Ligase PafA. *J. Am. Chem. Soc.* **135**, 6794–6797
